# TMEM106B C-terminal fragments drive nucleocytoplasmic transport failure and TDP-43 mislocalization in the aging human brain

**DOI:** 10.64898/2026.04.23.719939

**Authors:** Kedamawit Tilahun, Janani Parameswaran, Myles Dudley, Daniel Pun, Fuying Ma, Jessee Zhang, Tatiana Bold, Jie Jiang

## Abstract

TMEM106B is a lysosomal membrane protein and major genetic modifier of multiple neurodegenerative diseases, including frontotemporal lobar degeneration, Alzheimer’s disease, and amyotrophic lateral sclerosis. Proteolytically generated C-terminal fragments of TMEM106B assemble into amyloid fibrils that accumulate in the brains of individuals with neurodegenerative disease and in cognitively normal aged adults, yet how these fibrils produce neuronal dysfunction has remained unclear. Here, we show that cytosolic and lysosome-directed TMEM106B C-terminal fragments (CTF and gCTF) form detergent-insoluble amyloid aggregates, drive redistribution of endogenous TDP-43 from the nucleus to the cytoplasm, and accelerate neuronal death. Unbiased proximity proteomics identified the inner nuclear membrane LAP1-TorsinA axis as a fragment-specific interactome, and co-immunoprecipitation confirmed a direct physical interaction between gCTF and LAP1 that was not observed with full-length TMEM106B. Fragment expression disrupted Lamin B1 organization, mislocalized the nuclear import machinery KPNB1 and RanGAP1, and impaired importin-dependent nuclear transport in primary cortical neurons. Critically, neurons harboring endogenous TMEM106B fibrillar pathology in aged human frontal cortex exhibited the same phenotypes, namely disrupted Lamin B1 and LAP1 localization and cytoplasmic redistribution of TDP-43, whereas fibril-negative neurons from the same cases and younger control tissue retained intact nuclear envelope organization. These findings define TMEM106B proteinopathy as an upstream driver of nuclear envelope disruption and nucleocytoplasmic transport failure, linking a widespread feature of brain aging to a central mechanism of neurodegeneration.

## Introduction

Protein aggregation is a prominent feature of normal brain aging and a defining hallmark of many neurodegenerative diseases^1, 2^. Across disorders such as Alzheimer’s disease (AD), frontotemporal lobar degeneration (FTLD), amyotrophic lateral sclerosis (ALS), and Parkinson’s disease (PD), neurons accumulate insoluble protein assemblies that interfere with cellular homeostasis and contribute to neuronal vulnerability^3^. Although individual diseases are often defined by signature pathological proteins such as tau, amyloid β, α-synuclein, or TDP-43, neuropathological analyses increasingly reveal overlapping and co-existing protein aggregates within the same brains and neuronal populations^4–11^. Central to this mixed pathology is TDP-43, a normally nuclear RNA-binding protein whose cytoplasmic accumulation and nuclear depletion are defining features of ALS and FTLD. Notably, TDP-43 dysfunction is also a key player in the clinical features associated with AD and is frequently observed in older people without FTLD^12–14^. These observations suggest that neurodegeneration and brain aging are not driven by isolated proteinopathies but rather by convergent disruptions of shared cellular systems.

TMEM106B has emerged as a protein of growing interest at the intersection of aging and neurodegeneration. TMEM106B is a type II lysosomal membrane protein expressed throughout the central nervous system and implicated in regulating lysosomal morphology, trafficking, and function^15–18^. A genome-wide association study first identified common variants at 7p21 as risk factors specifically for the TDP-43 subtype of FTLD^19^, and subsequent neuropathological studies established that the TMEM106B risk genotype tracks with greater TDP-43 pathological burden, most prominently in granulin (GRN) mutation carriers, where risk alleles increase TDP-43 inclusion load and modify age at onset^20–22^. The genotype–TDP-43 relationship extends beyond FTLD to cognitive impairment in ALS^23^, with TDP-43 pathology in aging brains that lack an FTLD diagnosis^24^, and with hippocampal sclerosis of aging^25^, a distinct TDP-43 proteinopathy of the aging brain. Parallel work linked TMEM106B variants to AD and to cognitive dysfunction in Parkinson’s disease^26, 27^, collectively positioning TMEM106B as a genetic modifier of TDP-43 proteinopathies across aging and neurodegeneration.

Early studies proposed that TMEM106B risk SNPs confer pathogenicity by elevating its expression, and indeed, increased TMEM106B levels in cellular and mouse models produce enlarged lysosomes, disrupted endolysosomal trafficking, elevated lipofuscin accumulation, and impaired lysosomal exocytosis^15, 17, 18^. Yet loss of TMEM106B produces an equally disruptive phenotypic profile: Tmem106b knockout mice exhibit lysosomal trafficking defects, myelination abnormalities, synaptic loss, and progressive neurodegeneration^16, 28–30^. Most strikingly, Tmem106b deficiency severely exacerbates the *Grn* knockout phenotype, producing shortened lifespan, motor deficits, widespread microglial and autophagic activation, and prominent phosphorylated TDP-43 deposition that is not observed in single *Grn* knockouts^28, 29, 31^. This genetic interaction provides an *in vivo* link between TMEM106B, progranulin biology, and TDP-43 proteinopathy. These collective findings indicate that neuronal health requires precise homeostatic control of TMEM106B abundance, with both excess and insufficiency impairing lysosomal function.

An additional layer of TMEM106B biology has recently been revealed by the discovery that lysosomal proteases cleave full-length TMEM106B to generate a C-terminal fragment that assembles into β-sheet-rich amyloid fibrils^4, 5, 9, 32^. Cryo-EM studies demonstrated that these fibrils, rather than TDP-43 itself, constitute the previously unidentified amyloid present in FTLD-TDP brain^4, 9^, and that they accumulate across a broad spectrum of neurodegenerative diseases^5, 8^ as well as in cognitively normal individuals over age 50 in an age-dependent manner^4, 32^. The disease relevance of this fibril accumulation has been substantiated by pathological studies demonstrating that TMEM106B fibril core burden is strongly associated with the risk allele in a gene dosage-dependent manner and that increased fibril core deposition correlates robustly with enhanced TDP-43 dysfunction and disease severity in FTLD-TDP^11, 33^. Mechanistically, the T185 risk variant (rs3173615) has recently been shown to prolong the half-life of the fibril-forming fragment within lysosomes, culminating in lysosomal membrane rupture, phosphorylated TDP-43 inclusions, and neurodegeneration *in vivo*^33^. Despite this progress, the downstream cellular consequences of TMEM106B C-terminal fragment accumulation — particularly how fibril-forming fragments engage pathways beyond the lysosome to produce neuronal dysfunction — remain unknown.

Here, we investigate the cellular consequences of TMEM106B C-terminal fragment proteinopathy using complementary mammalian and neuronal model systems. We examined both a cytosolic C-terminal fragment (CTF) and a lysosome-directed, glycosylated C-terminal fragment (gCTF), and demonstrate that these fragments readily form detergent-insoluble, amyloid-like aggregates that resemble those found in patient tissues. Crucially, we find that the presence of TMEM106B fibrils is sufficient to drive the redistribution of endogenous TDP-43 from the nucleus to the cytoplasm and that neurons harboring endogenous TMEM106B fibrillar pathology in aged human frontal cortex show the same phenotype. Mechanistically, proximity-labeling proteomics identifies a fragment-specific interactome enriched for nuclear envelope components, including LAP1 and Torsin family members, whose disruption is accompanied by impaired nucleocytoplasmic transport and accelerated neuronal death. These results provide a mechanistic link between TMEM106B proteinopathy and the nuclear envelope defects that contribute to TDP-43 pathology in both normal aging and neurodegeneration.

## Results

### TMEM106B CTFs Form Amyloid Structures and Induce Neuronal Toxicity

The lysosomal TMEM106B protein consists of three major domains: an N-terminal cytosolic domain, a single transmembrane helix, and a C-terminal luminal domain that encompasses the recently identified fibril core associated with TMEM106B proteinopathy (**Figure 1A**) ^4, 5, 9, 32^. To dissect cellular consequences of TMEM106B fibril accumulation, we established constructs expressing either a cytosolic C-terminal fragment (CTF) or a lysosome-directed glycosylated CTF (gCTF) (**Figure 1A**). The gCTF construct was engineered by fusing the Progranulin (PGRN) signal peptide to the N-terminus and PGRN C-terminal sortilin-1 binding motif to the C-terminus to ensure ER/Golgi entry, luminal processing, and lysosomal routing^34^. These two constructs allowed us to probe whether fragment pathogenicity depends on cytosolic exposure or can emerge following authentic luminal processing and glycosylation, mimicking the endogenous proteolytic fragment observed in the human brain. Immunoblotting revealed that gCTF displayed a characteristic size shift following PNGase F treatment, confirming robust N-linked glycosylation, whereas the cytosolic CTF showed no mobility changes, as expected (**Figure 1B**). We next assessed the biochemical solubility of these fragments using sequential RIPA and urea fractionation of transfected cell lysates. TMEM106B-FL ran at its canonical ∼43 kDa monomeric weight, and when samples were prepared without heat denaturation, an additional ∼70 kDa band was detected, corresponding to the previously reported glycosylated homodimer^35^. TMEM106B-CTF and gCTF migrated at molecular weights consistent with those seen in human patient tissue, with unglycosylated CTF resolving at ∼15 kDa and gCTF with tags appearing at ∼37 kDa. Notably, both CTF and gCTF were strongly enriched in the RIPA insoluble, urea-soluble fraction unlike TMEM106B-FL, which remained largely RIPA soluble (**Figure 1C)**. These results support that TMEM106B-CTF and gCTF have intrinsic propensities to form detergent-insoluble aggregates.

**Figure 1.**
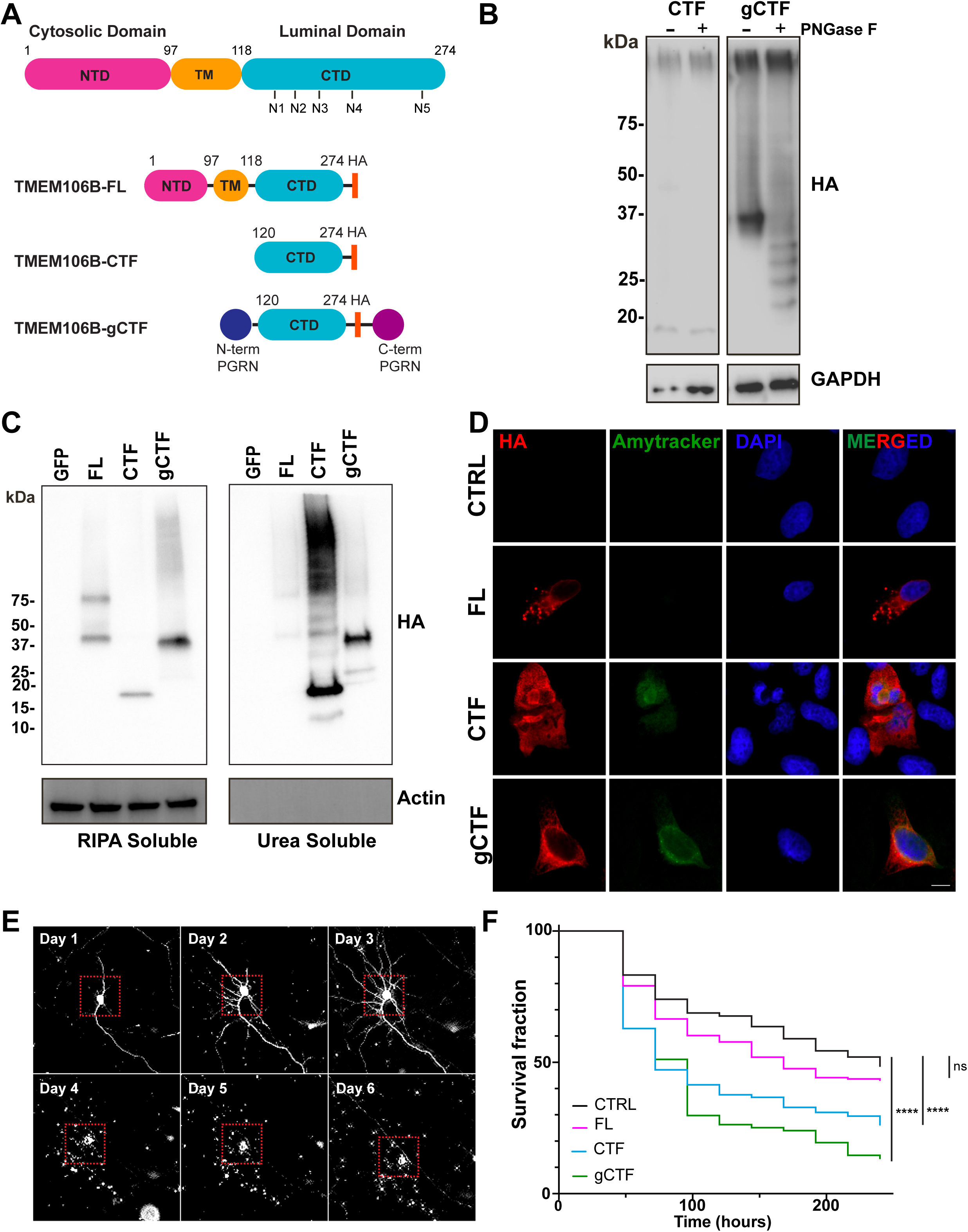
TMEM106B C-terminal fragments form insoluble amyloid aggregates and induce neuronal toxicity. **(A)** Schematic of TMEM106B domain architecture, showing the cytosolic N-terminal domain (NTD; residues 1–97), transmembrane domain (TM; residues 97–118), and luminal C-terminal domain (CTD; residues 118–274) with the five predicted N-linked glycosylation sites (N1–N5). Constructs used in this study are full-length TMEM106B (TMEM106B-FL), the cytosolic C-terminal fragment (TMEM106B-CTF, residues 120–274), and the lysosome-directed glycosylated C-terminal fragment (TMEM106B-gCTF), engineered by appending the progranulin (PGRN) signal peptide to the N-terminus to direct ER/Golgi entry, luminal processing, and glycosylation, and the PGRN C-terminal sortilin-1 binding motif to the C-terminus to enable lysosomal targeting. All constructs carry a C-terminal HA tag. **(B)** Anti-HA immunoblot of HA-tagged TMEM106B-CTF and gCTF with (+) or without (−) PNGase F treatment. **(C)** Anti-HA immunoblot of RIPA-soluble (left) and urea-soluble (right) fractions from U2OS cells expressing empty vector (GFP), TMEM106B-FL, CTF, or gCTF. Actin, loading control. **(D)** Representative immunofluorescence images of U2OS cells expressing empty vector (CTRL), TMEM106B-FL, CTF, or gCTF, stained for HA (red), Amytracker 540 (green; amyloid-binding dye), and DAPI (blue). Scale bar, 10 μm. **(E)** Representative longitudinal brightfield / mApple fluorescence time-lapse images of a primary cortical neuron expressing TMEM106B-gCTF and mApple, tracked every 24 h for 6 days. Dashed red boxes indicate the tracked cell body. **(F)** Kaplan–Meier survival curves for primary cortical neurons co-transfected at DIV4 with mApple and empty vector (CTRL), TMEM106B-FL, CTF, or gCTF, tracked by automated longitudinal microscopy over ∼240 h. Statistics: Cox proportional hazards regression relative to CTRL; n = 100 neurons per condition and experiments were repeated three times.

To determine whether these biochemically insoluble species formed amyloid-like structures, we performed immunofluorescence imaging with the amyloid-binding dye Amytracker 540. As observed in previous reports, Amytracker 540 labelled TMEM106B inclusions in frontal cortex of a 72-year-old human subject (**Figure S1A**)^10^, validating the dye’s detection of endogenous TMEM106B amyloid. Similarly, expressing CTF and gCTF in U2OS cells revealed robust amyloid aggregates whereas TMEM106B-FL produced no detectable amyloid signal (**Figure 1D**).

Given the prior links between elevated TMEM106B levels and lysosomal dysfunction, we next examined lysosomal morphology using the lysosomal marker LAMP1. While TMEM106B-FL expression in U2OS cells resulted in pronounced lysosomal expansion and a significant increase in total LAMP1 fluorescence intensity, TMEM106B-CTF or gCTF produced minimal effects on lysosomal morphology and did not significantly alter LAMP1 signal (**Figure S1B-C**). These results indicate that CTF- and gCTF-driven phenotypes are not a consequence of TMEM106B overexpression-associated lysosomal enlargement and instead reflect properties intrinsic to the fragments themselves.

Protein inclusions formed by several disease-related proteins have been shown to trigger cellular toxicity^3^. To test whether TMEM106B CTFs cause neuronal toxicity, we co-transfected TMEM106B-FL, CTF, or gCTF together with mApple in mouse primary cortical neurons at 4 days *in vitro* and used automated longitudinal microscopy to track the survival of hundreds of neurons marked by mApple fluorescence. Cell death was scored based on loss of fluorescence intensity, rounding of the cell body, and/or loss of neurite integrity, as exemplified by a representative gCTF-expressing neuron that underwent progressive neurite fragmentation starting at Day 4 (**Figure 1E**). Neurons expressing TMEM106B-CTF and gCTF exhibited an accelerated time-to-death relative to those expressing empty vector, whereas TMEM106B-FL caused only modest, statistically non-significant toxicity (**Figure 1F**). Together, these data demonstrate that TMEM106B C-terminal fragments induce neuronal toxicity.

### TMEM106B Pathology Correlates with TDP-43 Mislocalization

Prior human postmortem studies have reported that TMEM106B fibril deposition tracks with greater phosphorylated TDP-43 (pTDP-43) pathology, supporting a relationship between TMEM106B burden and TDP-43 proteinopathy^11, 36^. To dissect this relationship mechanistically, we first asked whether TMEM106B fragments induce canonical pTDP-43 pathology. A positive-control construct encoding C-terminal 25 kDa TDP-43 fragment (TDP-25) formed cytoplasmic puncta that robustly colocalized with pTDP-43 immunoreactivity, whereas none of the TMEM106B constructs, including TMEM106B-CTF and gCTF, induced detectable pTDP-43 (**Figure S2A**). Because nuclear depletion of TDP-43 is recognized as an early pathological event that precedes cytoplasmic pTDP-43 inclusion formation^37^, we next assessed whether TMEM106B fragments affect endogenous TDP-43 distribution. Both TMEM106B-CTF and gCTF drove redistribution of nuclear TDP-43 to the cytoplasm, whereas TMEM106B-FL had minimal effect (**Figure S2B**). We next asked whether TMEM106B C-terminal fragments are sufficient to drive TDP-43 mislocalization in a defined neuronal model. Neurons expressing TMEM106B-CTF or gCTF displayed redistribution of endogenous TDP-43 from the nucleus to the cytoplasm, whereas neurons expressing empty vector or TMEM106B-FL maintained strong TDP-43 nuclear enrichment **(Figure 2A-B).**

**Figure 2.**
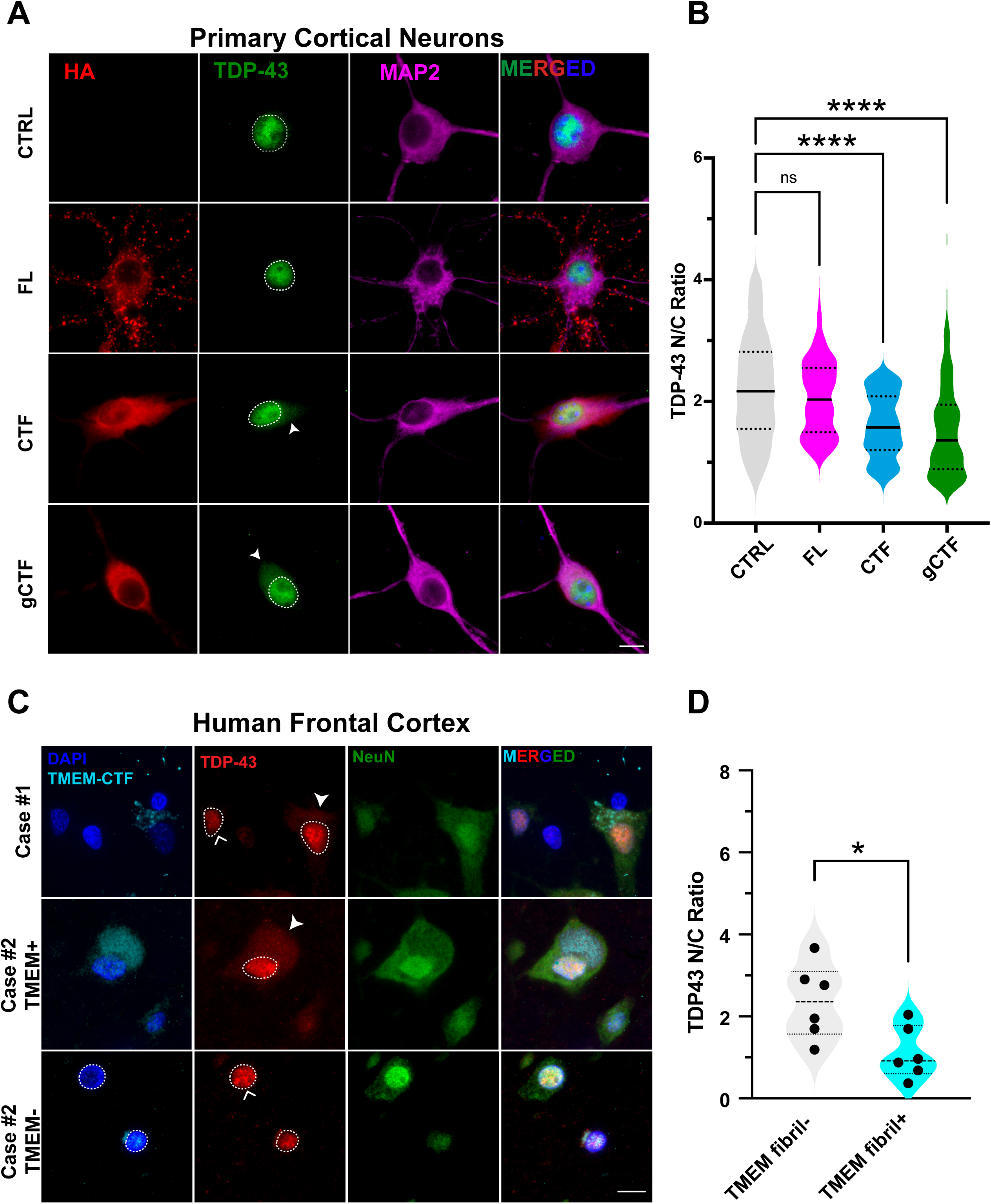
TMEM106B C-terminal fragments drive TDP-43 nuclear-to-cytoplasmic mislocalization in primary cortical neurons and in aged human frontal cortex. **(A)** Representative immunofluorescence images of primary cortical neurons transduced at DIV4 with AAV expressing empty vector (CTRL), TMEM106B-FL, CTF, or gCTF, stained for HA (red), TDP-43 (green), and MAP2 (magenta, merged). Dashed outlines denote nuclei; arrowheads indicate cytoplasmic TDP-43 signal in CTF-and gCTF-expressing neurons. **(B)** Quantification of TDP-43 nuclear-to-cytoplasmic (N/C) ratio in primary cortical neurons. Violin plots show median (solid line) and interquartile range (dashed lines). ****p < 0.0001; ns, not significant (one-way ANOVA with Tukey’s post-hoc test); n = 40-50 neurons per condition across 3 independent cultures. **(C)** Representative immunofluorescence images of postmortem human frontal cortex from aged, neurologically normal individuals (Case #1, Case #2 TMEM+ and TMEM− neurons shown), stained with DAPI and anti-TMEM106B luminal domain (TMEM-CTF; cyan/blue), TDP-43 (red), and NeuN (green). Dashed outlines denote neuronal nuclei; filled arrowheads indicate cytoplasmic TDP-43 in fibril-positive neurons; open arrowheads highlight nuclear TDP-43 retention in fibril-negative neurons within the same tissue. **(D)** Quantification of TDP-43 N/C ratio in NeuN+ neurons from six aged control individuals, classified as TMEM fibril-negative (TMEM fibril−) or fibril-positive (TMEM fibril+) within the same case. Each dot represents the mean value per case (n = 6 cases per group). Violin plots show median and interquartile range. *p < 0.05 (paired two-tailed t-test). Scale bar, 10 μm.

To determine whether the same association holds in aged human brain, we analyzed postmortem frontal cortex from six aged, neurologically normal human individuals using a luminal domain-specific antibody previously validated to recognize the proteolytically generated, fibril-forming C-terminal fragment in human autopsy tissue^38^. This antibody enabled us to distinguish NeuN-positive neurons with detectable TMEM106B fibrils from neighboring fibril-negative neurons within the same case. Across all individuals with detectable fibrillar pathology, neurons containing TMEM106B fibrils showed loss of nuclear TDP-43 with cytoplasmic redistribution, whereas adjacent fibril-negative neurons retained strong nuclear TDP-43 enrichment **(Figure 2C and S3A).** Notably, one 22-year-old young human case did not exhibit any TMEM106B fibrillar pathology, and no TDP-43 mislocalization was observed in that individual **(Figure S3B).** Quantification confirmed a significantly reduced TDP-43 nuclear-to-cytoplasmic ratio in fibril-positive neurons compared with fibril-negative neurons from the same cases **(Figure 2D).** Together, these findings establish that TMEM106B fibrillar pathology is associated with TDP-43 mislocalization in aged human cortex, and that fragment accumulation alone is sufficient to drive this phenotype in primary neurons.

### TMEM106B Fragments Disrupt Nuclear Envelope Integrity and Nucleocytoplasmic Transport

Having established that TMEM106B C-terminal fragments drive TDP-43 mislocalization in neuronal models, we next sought to understand the underlying mechanism. TDP-43 is a predominantly nuclear protein whose cytoplasmic accumulation in disease has been linked to impaired nuclear import^39^, but the upstream events that precipitate this mislocalization remain poorly defined. An initial clue came from U2OS cells, where many TMEM106B-CTF-expressing cells displayed abnormal nuclear indentations and reduced nuclear circularity, a phenotype not observed with TMEM106B-FL or gCTF (**Figure S4A -S4B**). This nuclear shape distortion was not recapitulated in primary cortical neurons transduced with AAV-CTF or AAV-gCTF (**Figure S4C-S4D**), indicating that the magnitude of structural deformation depends on cell type and/or expression level. Nonetheless, the appearance of nuclear shape abnormalities was striking given the typically stable nuclear morphology of these cells, prompting us to examine nuclear envelope integrity directly.

Prior studies have shown that protein aggregates can compromise nuclear envelope integrity, manifested as disrupted Lamin B1 organization at the nuclear rim^40–42^. We therefore examined Lamin B1 distribution in primary cortical neurons expressing TMEM106B constructs. Neurons expressing CTF or gCTF exhibited significantly reduced Lamin B1 intensity at the nuclear rim compared with control or FL-expressing neurons, despite unchanged overall nuclear shape (**Figure 3A and B**). This phenotype was recapitulated in U2OS cells, where CTF and gCTF expression similarly reduced Lamin B1 rim intensity relative to control and FL conditions (**Figure S4E-F**), indicating that nuclear lamina disruption is a consistent consequence of fragment expression across both cell types tested.

**Figure 3.**
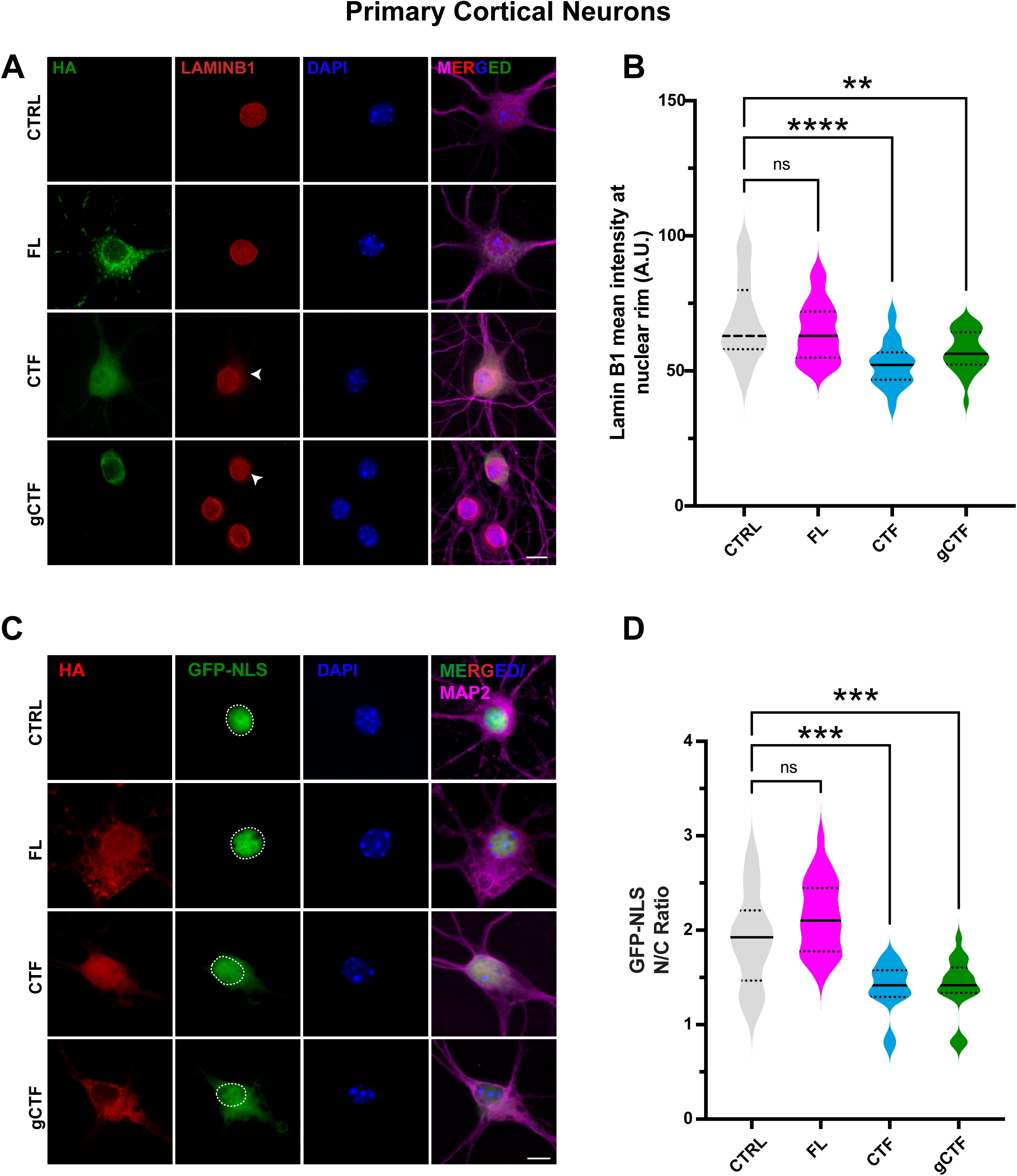
TMEM106B C-terminal fragments disrupt Lamin B1 organization and impair nuclear import in primary cortical neurons. **(A)** Representative immunofluorescence images of primary cortical neurons transduced at DIV4 with AAV expressing empty vector (CTRL), TMEM106B-FL, CTF, or gCTF, stained for HA (green), Lamin B1 (red), and DAPI (blue), with MAP2 shown in the merged panel (magenta). Arrowheads indicate reduced Lamin B1 signal at the nuclear rim in CTF- and gCTF-expressing neurons. **(B)** Quantification of Lamin B1 mean intensity at the nuclear rim. Violin plots show median and interquartile range. ****p < 0.0001; **p < 0.01; ns, not significant (one-way ANOVA with Tukey’s post-hoc test); n = 40 –50 neurons per condition across 3 independent cultures. **(C)** Representative immunofluorescence images of primary cortical neurons co-transduced with AAV-GFP-NLS (nuclear import reporter; green) and HA-tagged CTRL, TMEM106B-FL, CTF, or gCTF (red), counterstained with DAPI (blue) and MAP2 (magenta, merged). Dashed outlines denote nuclei; arrowheads highlight cytoplasmic GFP-NLS accumulation in CTF- and gCTF-expressing neurons. **(D)** Quantification of GFP-NLS N/C ratio. Violin plots show median and interquartile range. ****p < 0.0001; ***p < 0.001; **p < 0.01; ns, not significant (one-way ANOVA with Tukey’s post-hoc test); n = 30-40 neurons per condition across 3 independent cultures. Scale bar, 10 μm.

The structural disruption of the nuclear lamina raised the question of whether fragment expression also perturbs the molecular machinery of nucleocytoplasmic transport. The nuclear import receptor KPNB1 and the Ran GTPase regulator RanGAP1 are essential for maintaining the gradient that drives importin-β-dependent transport. In U2OS cells expressing TMEM106B-CTF or gCTF, both KPNB1 and RanGAP appeared qualitatively disrupted in their subcellular distributions, including apparent loss of normal perinuclear enrichment and aberrant cytoplasmic or diffuse localization (**Figure S4G-H**). These alterations are consistent with impaired organization of the nuclear import apparatus downstream of nuclear envelope disruption.

To determine whether these structural perturbations translate to a functional deficit, we employed a GFP-NLS reporter that provides a sensitive readout of importin-dependent trafficking. Similar NLS-based reporters have been widely used to detect nuclear import impairments in aggregate-bearing cells across multiple neurodegeneration models, including systems in which TDP-43 mislocalization is driven by transport defects^43–45^. Primary cortical neurons co-transduced with GFP-NLS and CTF or gCTF displayed prominent cytoplasmic accumulation of the reporter, whereas control and FL-expressing neurons retained strong nuclear GFP-NLS signal (**Figure 3C**). Quantification of the nuclear-to-cytoplasmic GFP-NLS ratio confirmed a significant reduction in both fragment-expressing conditions (**Figure 3D**). Together, the reduced GFP-NLS nuclear import, disorganization of KPNB1 and RanGAP1, and Lamin B1 depletion from the nuclear rim support a coherent structural-to-functional failure pathway in neurons, in which TMEM106B fragment accumulation compromises the nuclear lamina and, with it, the transport machinery required to maintain TDP-43 nuclear localization.

### Proximity proteomics identifies the LAP1-Torsin axis as a TMEM106B gCTF interactor

The structural and functional nuclear envelope defects observed in fragment-expressing neurons raised a mechanistic question: how does a lysosomally-generated fragment engage the nuclear envelope machinery? To gain unbiased mechanistic insight into this question, we employed an APEX2-based proximity labeling strategy in SH-SY5Y neuroblastoma cells using TMEM106B-gCTF fused to a V5-APEX2 tag. We confirmed that the fusion retained biological activity, as gCTF-V5-APEX2-expressing cells displayed a significant reduction in TDP-43 nuclear-to-cytoplasmic ratio compared to V5-APEX2 controls (**Figure 4A-B**), and streptavidin blotting confirmed efficient proximity labeling (**Figure 4C**). Mass spectrometry identified 77 proteins significantly enriched in the gCTF proximal proteome (**Figure 4D**; **Supplementary Table 2**). Gene Ontology analysis revealed expected enrichment of ER stress, protein folding, and glycoprotein metabolic terms, consistent with the luminal biogenesis route of gCTF, alongside significant enrichment of nucleocytoplasmic transport and nuclear pore complex assembly terms (**Figure S5**). These NE-associated terms included TorsinA (TOR1A), Torsin-1A interacting protein 1 (TOR1AIP1, which encodes Lamina-Associated Polypeptide 1 [LAP1]), and NUP210, among their constituent proteins, nominating the LAP1-TorsinA axis as a candidate TMEM106B gCTF-interacting complex. To validate the LAP1 interaction, we co-transfected V5-tagged LAP1 with HA-tagged TMEM106B-FL or gCTF and performed co-immunoprecipitation via HA pulldown. V5-LAP1 was recovered specifically in the gCTF immunoprecipitate and not in FL or control conditions (**Figure 4E**), establishing that LAP1 engages the cleaved C-terminal fragment but not full-length TMEM106B.

**Figure 4.**
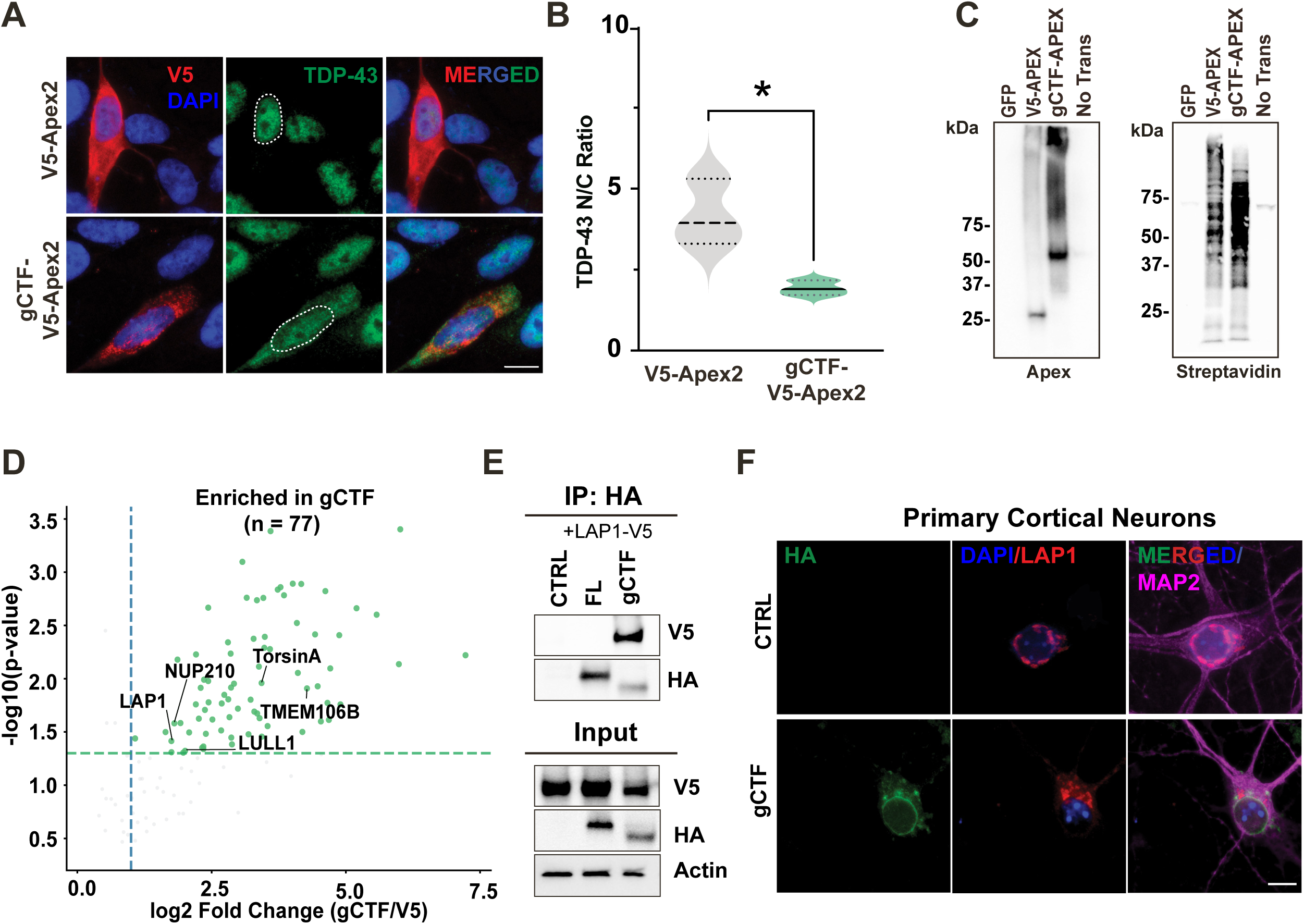
Proximity proteomics identifies the LAP1–TorsinA axis as a TMEM106B gCTF interactor. **(A)** Representative immunofluorescence images of SH-SY5Y cells expressing V5-APEX2 (CTRL) or TMEM106B-gCTF-V5-APEX2 (gCTF), stained for V5 (red), TDP-43 (green), and DAPI (blue). Dashed outlines denote nuclei. Scale bar, 10 μm. **(B)** Quantification of TDP-43 N/C ratio in SH-SY5Y cells expressing V5-APEX2 (V5) or gCTF-APEX2 (gCTF). *p < 0.05 (unpaired two-tailed t-test). **(C)** Anti-APEX2 (left) and streptavidin (right) immunoblots of lysates from SH-SY5Y cells expressing GFP, V5-APEX2, gCTF-APEX2, or untransfected control (No Trans), treated with biotin-phenol and H₂O₂. Streptavidin signal confirms efficient biotinylation of proximal proteins in V5-APEX2 and gCTF-APEX2 conditions. **(D)** Volcano plot of mass spectrometry results comparing the gCTF-APEX2 proximal proteome to V5-APEX2 controls. Green dots indicate significantly enriched proteins (log₂ fold-change > 1, p < 0.05; dashed thresholds). Highlighted hits include LAP1 (TOR1AIP1), TorsinA (TOR1A), LULL1 (TOR1AIP2), NUP210, and TMEM106B itself. **(E)** HA co-immunoprecipitation of V5-LAP1 with HA-tagged TMEM106B-FL, gCTF, or empty vector (CTRL) in co-transfected U2OS cells. **(F)** Representative immunofluorescence images of primary cortical neurons expressing HA-tagged empty vector (CTRL) or gCTF, stained for HA (green), LAP1 (red), DAPI (blue), and MAP2 (magenta, merged). Scale bar, 10 μm.

Because LAP1 is an inner nuclear membrane protein that organizes TorsinA-dependent nuclear pore complex (NPC) maturation and nuclear lamina homeostasis, we next asked whether LAP1 distribution is perturbed in TMEM106B fragment-expressing neurons. In primary cortical neurons, LAP1 was enriched at the nuclear rim in control neurons. In contrast, gCTF-expressing neurons displayed LAP1 mislocalization away from the nuclear envelope (**Figure 4F**). Together, these data establish that gCTF physically engages LAP1 and disrupts its organization at the inner nuclear membrane, providing a molecular interaction that links TMEM106B fragment accumulation to the nuclear envelope defects we observed.

### Nuclear envelope defects in endogenous TMEM106B fibril-positive neurons in aged human cortex

To determine whether the nuclear envelope defects observed in our cell-based and primary neuron models occur in neurons harboring endogenous TMEM106B fibrillar pathology, we examined Lamin B1 and LAP1 organization in aged human frontal cortex. Using the luminal domain-specific antibody described above to distinguish TMEM106B fibril-positive from fibril-negative neurons in the same tissue, we assessed nuclear envelope integrity in both populations across six aged, neurologically normal individuals.

Fibril-positive neurons consistently displayed aberrant, fragmented, or mislocalized Lamin B1 signal at the nuclear periphery, whereas adjacent fibril-negative neurons exhibited smooth, continuous Lamin B1 staining along the nuclear rim (**Figure 5A, S6A**). TOM20 staining in the same fibril-positive neurons revealed intact cytoplasmic mitochondrial signal, indicating that Lamin B1 disruption is specific to the nuclear envelope and not a reflection of generalized cellular disorganization **(Figure S7).** Frontal cortex from a young individual lacking detectable TMEM106B fibrillar pathology (Case #7, age 22) displayed uniformly smooth, continuous Lamin B1 staining with no evidence of nuclear envelope disruption (**Figure S6C**), establishing that the phenotype is specifically coupled to fibril accumulation rather than to aging or postmortem artifact.

**Figure 5.**
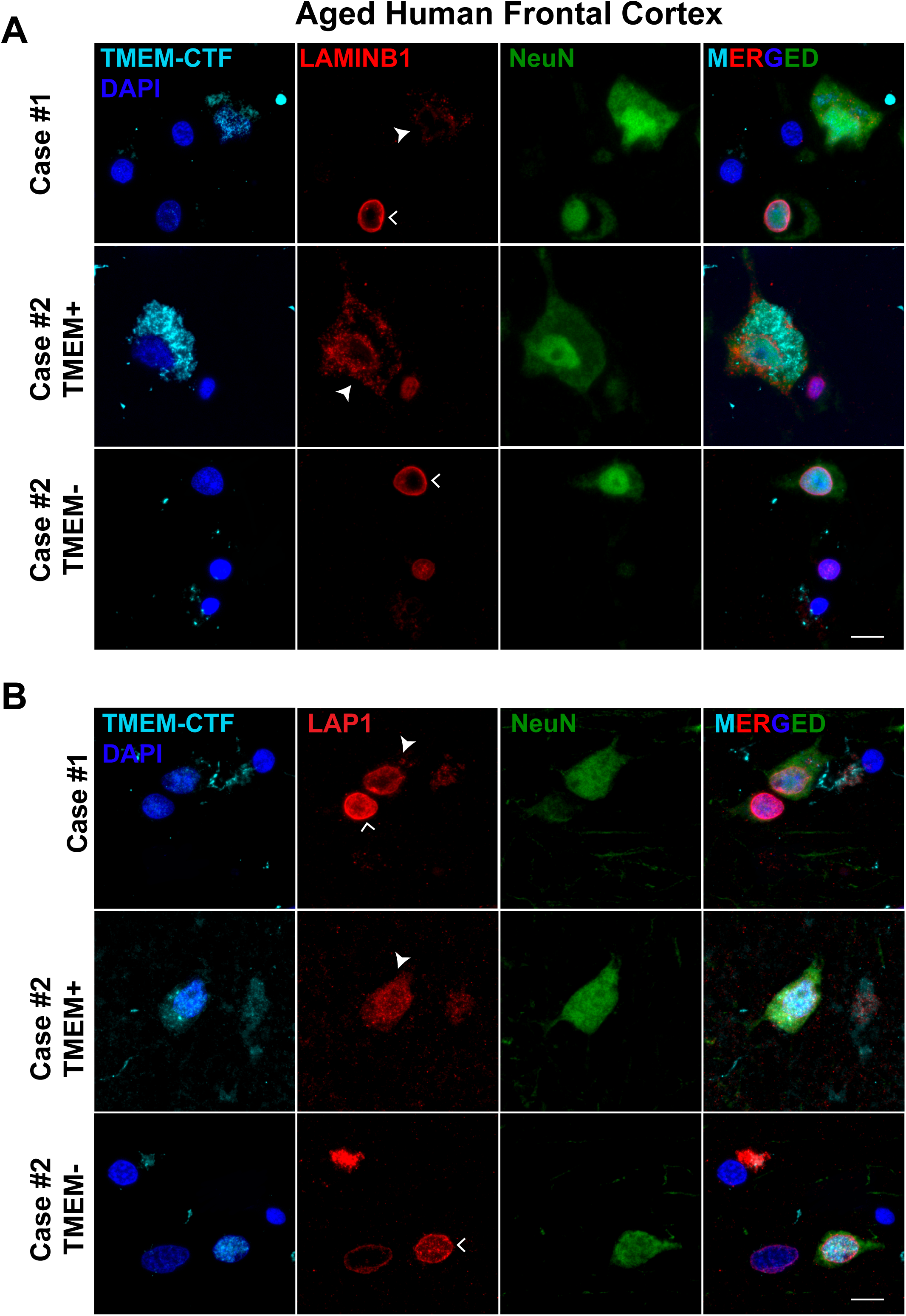
Endogenous TMEM106B fibrillar pathology is associated with disrupted Lamin B1 and LAP1 organization in aged human frontal cortex. **(A)** Representative immunofluorescence images of postmortem frontal cortex from aged, neurologically normal individuals, stained with DAPI and anti-TMEM106B luminal domain (TMEM-CTF; cyan/blue), Lamin B1 (red), and NeuN (green). Cases shown: Case #1 (fibril-positive neuron, top), Case #2 TMEM+ (fibril-positive, middle), and Case #2 TMEM− (fibril-negative neuron from the same case, bottom). Filled arrowheads fibril-positive neurons; open arrowheads indicate fibril-negative neurons. **(B)** Representative immunofluorescence images of the same cases, stained for TMEM-CTF/DAPI (cyan/blue), LAP1 (red), and NeuN (green). Fibril-positive neurons display irregular or fragmented LAP1 staining at the nuclear rim (filled arrowheads), whereas fibril-negative neurons retain continuous LAP1 localization (open arrowheads). Scale bar, 10 μm.

We next examined LAP1 organization in the same cohort. Fibril-positive neurons consistently displayed irregular or mislocalized LAP1 staining, whereas neighboring fibril-negative neurons retained continuous LAP1 localization at the nuclear rim (**Figure 5B, S6B**). The young fibril-negative case likewise showed uniformly intact LAP1 organization **(Figure S6C),** supporting a fibril-dependent disruption.

Together, these findings demonstrate that Lamin B1 and LAP1 disorganization are consistent features of TMEM106B fibril-positive neurons in aged human cortex, linking the fragment-specific interactome and structural nuclear envelope defects identified in our models to endogenous TMEM106B pathology *in vivo*.

## Discussion

In this study, we examined the cellular consequences of TMEM106B C-terminal fragment accumulation using complementary cellular and primary neuron models alongside aged human frontal cortex. We show that cytosolic and lysosome-directed fragments (CTF and gCTF) readily assemble into detergent-insoluble, amyloid-like aggregates, drive redistribution of endogenous TDP-43 from the nucleus to the cytoplasm, disrupt nuclear envelope organization and the nucleocytoplasmic transport machinery, and accelerate neuronal death. Unbiased proximity proteomics identified the inner nuclear membrane LAP1-Torsin axis as a fragment-specific interactome, and we validated both physical interaction with LAP1 by co-immunoprecipitation and altered LAP1 and Lamin B1 organization in TMEM106B fibril-positive neurons across six aged human individuals. These results define a mechanistically separable arm of TMEM106B pathobiology, distinct from lysosomal dysfunction, and place TMEM106B proteinopathy upstream of the nuclear transport failure that characterizes multiple neurodegenerative diseases.

A notable feature of our cellular models is that neither CTF nor gCTF expression produced detectable phosphorylated TDP-43 inclusions, despite the strong genetic association between TMEM106B risk variants and pTDP-43 burden in FTLD-TDP^11, 24, 33, 46^. Instead, both fragments consistently drove redistribution of endogenous TDP-43 from the nucleus to the cytoplasm. This dissociation is mechanistically informative rather than paradoxical. Extensive evidence from ALS and FTD studies supports the view that nuclear depletion of TDP-43 is an early event that precedes and contributes to the formation of cytoplasmic phosphorylated inclusions: pathological analysis of ALS spinal cord has shown that diffuse cytoplasmic TDP-43 mislocalization is observed in neurons that retain residual nuclear TDP-43 signal and that these early mislocalized species are phosphorylated but poorly ubiquitylated, with compact skein-like and ubiquitin-positive inclusions emerging only as nuclear TDP-43 becomes fully depleted^37^. Consistent with a sequential model, monomerization of TDP-43 has been identified as an early pathological event that drives nuclear export via Nxf1-dependent mechanisms, with cytoplasmic accumulation then promoting phosphorylation and aggregation as downstream consequences^47^. Supporting our observation, a recent study examining endogenous TMEM106B pathology in C9orf72-ALS mouse models and matched human C9-ALS and ALS/FTD postmortem tissue found that neurons bearing TMEM106B cytoplasmic puncta had a significantly reduced TDP-43 nuclear-to-cytoplasmic ratio compared to TMEM106B-negative neurons in the same sections, establishing a cell-autonomous correlation between TMEM106B pathology and TDP-43 nuclear clearance in a disease context^48^. Taken together, these observations suggest that TMEM106B CTF-driven nuclear transport impairment represents an upstream mechanism that primes neurons toward TDP-43 mislocalization, and that the progression from this early mislocalization state to full pTDP-43 proteinopathy likely requires convergent stressors, including the lysosomal damage and fibril-induced seeding described by companion studies, to cross the threshold into irreversible aggregation^33, 49^.

Nucleocytoplasmic transport (NCT) failure has emerged over the past decade as a converging mechanism across ALS, FTD, and related neurodegenerative diseases^45, 50–53^. Seminal work in C9orf72-ALS identified RanGAP sequestration by G4C2 repeat RNA foci as a cause of Ran gradient collapse and TDP-43 nuclear depletion^45^, and unbiased genetic screens in Drosophila independently identified components of the nuclear pore complex and Ran cycle as potent modifiers of repeat-mediated neurodegeneration^50^. Pathological TDP-43 aggregates themselves reciprocally disrupt NPC morphology and NCT, establishing a feedforward loop in which transport failure and proteinopathy mutually amplify each other^52^. KPNB1 deficits are sufficient to induce TDP-43 mislocalization^39^, while enhancing KPNB1 and related nuclear import receptor availability dissolves pathological TDP-43 assemblies and restores nuclear localization across multiple ALS/FTD neuronal systems^54^. Despite this broad framework, the structural basis for how NCT failure is initiated upstream of TDP-43 proteinopathy, particularly the nuclear envelope scaffolding events that establish the preconditions for import, has remained poorly defined. Our findings identify TMEM106B CTFs as an upstream driver of this cascade, revealing their interaction with the nuclear envelope machinery that directly governs NPC integrity and providing a mechanistic link between CTF accumulation and the initiation of NCT failure.

Our APEX-based proximity proteomics of the glycosylated CTF recovered a selective enrichment of TorsinA (TOR1A), an AAA+ ATPase that is enzymatically inactive without cofactor engagement, alongside both of its known activators: LULL1 (TOR1AIP2), which stimulates TorsinA throughout the ER lumen, and LAP1 (TOR1AIP1), which activates TorsinA exclusively at the inner nuclear membrane^55, 56^. The co-recovery of TorsinA with a spatially distinct pair of activators, one ER-resident and one nuclear envelope-resident, raised the possibility that gCTF engages this axis at the nuclear envelope specifically — a possibility directly supported by co-immunoprecipitation demonstrating a physical interaction between gCTF and LAP1, but not full-length TMEM106B. The transmembrane nucleoporin NUP210, whose mislocalization is a structural hallmark of TorsinA-deficient neurons, was also enriched in the gCTF proximal proteome, further implicating the LAP1-TorsinA-NPC axis as the relevant interaction network. Supporting this nuclear-envelope connection, an independent TMEM106B fibril-interactor dataset likewise recovered canonical NE proteins like lamin A/C (LMNA), emerin (EMD), and nesprin-1 (SYNE1) along with nuclear-transport regulators such as RAN, KPNB1, and IPO5, further reinforcing that TMEM106B fibrils interface with nuclear-envelope-associated pathways^11^. LAP1’s luminal domain directly stimulates TorsinA AAA+ ATPase activity, and imbalanced LAP1 expression alone is sufficient to disrupt post-mitotic nuclear architecture^57^. Loss of TorsinA, whose dysfunction causes the neurodevelopmental movement disorder DYT1 dystonia, selectively disrupts the neuronal nuclear envelope^58^, and subsequent work established that this manifests as mislocalized NPC clusters lacking the late-recruited cytoplasmic nucleoporin Nup358 that persists into adulthood^59^, a consequence of excessive inner nuclear membrane budding at sites of nascent NPC assembly and delayed NPC formation in the absence of TorsinA^60^. Furthermore, TorsinA dysfunction causes abnormalities in Ran localization and reduced Lamin A/C intensity at the nuclear rim, phenotypes strikingly similar to those we observe in CTF-expressing neurons^59^. Consistent with these mechanistic relationships, gCTF expression disrupted LAP1 organization in primary cortical neurons, and a similar morphology was evident in TMEM106B fibril-positive neurons in human frontal cortex. While prior work has established this pathway in the context of genetic TorsinA loss during neurodevelopment, our findings identify TMEM106B CTF accumulation as an acquired mechanism that engages the LAP1-TorsinA axis in post-mitotic neurons, suggesting a convergent route to nuclear envelope dysfunction in age-associated proteinopathy.

CTF and gCTF expression also disrupted the subcellular organization of Lamin B1, KPNB1, and RanGAP1, components essential for nuclear lamina integrity and for maintaining the Ran-GTP/GDP gradient that drives importin-β-dependent nuclear import. Lamin B1 is known to decline from the nuclear rim in aging neural stem cells and neurons^61, 62^ and in ALS/FTD motor cortex independently of C9orf72 repeat expansions^63^ while Lamin A/C intensity is reduced in neurons with TorsinA dysfunction^59^, consistent with the nucleoplasmic domain of LAP1 directly contacting nuclear lamins^64^. The Lamin B1 reduction and disorganization we observe in CTF-expressing neurons and fibril-positive human cortex is consistent with an aging-related nuclear lamina vulnerability documented across neurodegeneration and aging contexts, and while the LAP1-TorsinA axis preferentially disrupts Lamin A/C rather than Lamin B1, the convergence on lamina instability across these distinct mechanisms further supports CTF-driven nuclear envelope structural failure as a biologically significant event. Critically, these structural perturbations translated into functional NCT impairment: using GFP-NLS reporter assays in primary cortical neurons, we directly demonstrated that CTF and gCTF expression increased cytoplasmic retention of the NLS-tagged reporter, confirming that TMEM106B fragments are sufficient to impair nuclear import in neurons. Import deficits have been reported across multiple ALS/FTD paradigms — including C9orf72 repeat toxicity^45, 50^, NPC injury models^52^, and arginine-rich dipeptide repeat expression^65^ — and blocking nuclear import is sufficient to cause neurotoxicity in neuronal systems^39, 66^. Our data now add TMEM106B CTF accumulation to this set of pathological triggers. The convergence of LAP1 disruption, Lamin B1, KPNB1 and RanGAP disorganization, and functional NLS reporter impairment in the same model system argues that these are not independent observations but sequential steps in a coherent structural-to-functional failure pathway. We propose CTFs engage the LAP1-TorsinA axis at the inner nuclear membrane, disrupt its role in NPC maturation and laminar organization, destabilize the Ran gradient, impair importin-β-dependent import, and ultimately drive nuclear depletion of TDP-43. Consistent with previous *in vivo* data showing that worms and mice expressing TMEM106B CTF exhibit age-dependent neurodegeneration^33, 67^, our neuronal models demonstrate that CTF and gCTF expression cause neuronal toxicity, indicating that these nuclear transport deficits are not bystander events but are associated with neuronal vulnerability.

An important caveat of our human tissue analyses concerns the interpretation of pathological findings in neurologically normal aged individuals. The presence of TMEM106B amyloid filaments in the brains of cognitively intact older adults, confirmed by cryo-EM and neuropathology in individuals without neurological diagnosis^4^, raises the question of whether the nuclear envelope disruption, Lamin B1 disorganization, and TDP-43 redistribution we observe in fibril-positive neurons represent genuinely pathological events or simply tolerated features of normal cellular aging. Several lines of evidence argue against the latter interpretation. First, these phenotypes are strictly age-gated and fibril-associated: they are entirely absent in younger control tissue and emerge in aged tissue in a pattern that tracks with TMEM106B fibril burden, indicating they are not a nonspecific consequence of chronological aging but are coupled to fibril accumulation. Second, the biological significance of fibril burden in ostensibly healthy brains is supported by population genetics: TMEM106B risk variants are associated with accelerated biological aging of the frontal cortex, reduced neuronal proportions, worse cognitive performance, and neuroinflammatory transcriptomic signatures even in the absence of diagnosed neurological disease, and these associations are selective for older individuals and absent in younger cohorts^68^. Third, and most compellingly, protective alleles at both TMEM106B and GRN are among the most strongly enriched variants in cognitively healthy centenarians relative to age-matched controls and Alzheimer’s disease cases^69^, and these are the very same alleles shown to reduce CTF accumulation and limit fibril formation^33, 70^. This genetic architecture implies that the molecular events downstream of CTF accumulation, including the nuclear envelope and nucleocytoplasmic transport defects we document, are not neutral aging byproducts but contribute meaningfully to the trajectory from cognitive resilience to vulnerability.

In summary, we identify TMEM106B C-terminal fragments as aggregation-prone species that disrupt nuclear envelope organization, impair nucleocytoplasmic transport, and drive TDP-43 mislocalization, phenotypes documented in cellular models, primary neurons, and human brain tissue spanning aging and neurodegeneration. These results define a mechanistically separable arm of TMEM106B pathobiology, distinct from lysosomal dysfunction and position TMEM106B proteinopathy as an upstream contributor to the nuclear transport failure that characterizes multiple neurodegenerative diseases.

### Limitations of the study and future directions

This study has several limitations that should be considered when interpreting our findings. First, our fragment-based expression systems which include cytosolic CTF and lysosome-directed gCTF constructs recapitulate key biochemical and morphological features of endogenous TMEM106B fragment pathology but may not fully reproduce the stoichiometry, processing kinetics, or subcellular context of CTF generation in the aging human brain. Endogenous CTF accumulation occurs gradually over decades within the lysosomal lumen, whereas our overexpression paradigms achieve supraphysiological fragment levels acutely, which may amplify or distort certain downstream phenotypes. Second, while APEX-based proximity labeling identified nuclear envelope components in the local environment of gCTF and co-immunoprecipitation supported a physical interaction with LAP1, the functional significance of the CTF-LAP1 interaction — including whether it is sufficient to impair TorsinA ATPase activity or NPC maturation *in vivo* — remains to be established. Third, our human tissue analyses, demonstrating consistent associations between TMEM106B fibril positivity and nuclear envelope disruption, Lamin B1 and LAP1 disorganization, and TDP-43 mislocalization, are correlative by nature. They do not establish causal relationships between CTF aggregation, nuclear envelope dysfunction, and neuronal degeneration, and the possibility that these phenotypes co-occur as parallel consequences of a shared upstream stress rather than a linear pathway cannot be excluded from postmortem data alone. Fourth, the identity of the specific lysosomal or cellular trigger that initiates CTF release from the lysosomal lumen into compartments proximal to the nuclear envelope remains unknown, and whether this requires lysosomal membrane rupture, as suggested by the recent cryo-ET data^33^, or proceeds through alternative trafficking routes under more physiological conditions is an important unresolved question. Future studies should determine whether genetic or pharmacological restoration of nuclear envelope integrity — through modulation of the LAP1-TorsinA axis — or enhancement of nucleocytoplasmic transport capacity can rescue CTF-induced phenotypes and whether such interventions modify TDP-43 localization and neuronal survival *in vivo*. It will also be important to define the endogenous timing, brain-region specificity, and cellular context of CTF generation and aggregation across the aging and disease trajectory, and to determine at what stage of fibril burden nuclear transport deficits become functionally consequential. Addressing these questions will be essential to establish whether TMEM106B-driven nuclear transport failure is a causal contributor to neurodegeneration or a biomarker of neuronal stress, and to evaluate its potential as a therapeutic target.

## Materials and Methods

### DNA constructs

Coding sequences for full-length human TMEM106B (TMEM106B-FL; amino acids 1–274, carrying the FTLD risk-associated T185 variant), a cytosolic C-terminal fragment (CTF; amino acids 120-274), and a glycosylated C-terminal fragment (gCTF) bearing an N-terminal PGRN signal peptide and a C-terminal Sortilin-1 binding motif were synthesized as gene blocks (Azenta Life Sciences). A gene block encoding the C-terminal 25 kDa fragment of TDP-43 (TDP-25; amino acids 208-414) was similarly synthesized and used as a positive control for pTDP-43 staining. Constructs were assembled into the pGW1 backbone with a C-terminal HA tag using Gibson Assembly (New England Biolabs). An empty pGW1 vector was used as a control. For longitudinal fluorescence microscopy, pGW1-mApple was used as a morphological marker. For AAV-mediated neuronal transduction, TMEM106B-FL, CTF, and gCTF were subcloned into the CTR4 AAV transfer vector and packaged into AAV8 capsids using the pDP8 Rep/Cap/Helper system. For APEX2-based proximity labeling, V5-APEX2 was PCR-amplified from pcDNA3.1_APEX2 and assembled into the CTR4 AAV backbone by Gibson Assembly; TMEM106B-gCTF was subsequently cloned into this vector to generate TMEM106B-gCTF-V5-APEX2. A GFP-NLS nuclear import reporter (Addgene, #104061) was packaged into AAV-PHP.eB. All plasmids were propagated in DH5α Escherichia coli and verified by Sanger sequencing.

### Cell culture and transfection

Human osteosarcoma (U2OS) cells were cultured in high-glucose DMEM (Invitrogen) supplemented with 10% fetal bovine serum (Corning), 4 mM GlutaMAX (Invitrogen), 100 U/mL penicillin, 100 μg/mL streptomycin, and 1% non-essential amino acids. Human neuroblastoma (SH-SY5Y) cells were cultured in DMEM/F-12 (Invitrogen) with identical supplements. Cells were maintained at 37°C in a humidified incubator with 5% CO₂. Transient transfections were performed using polyethylenimine (PEI, 25 kDa linear; Polysciences) at a DNA:PEI ratio of 1:3 (w/w) in Opti-MEM (Invitrogen). For immunofluorescence, cells were fixed 24 h after transfection. For immunoblotting, cell lysates were collected 48 h after transfection.

### Primary cortical neuronal culture and transfection/transduction

All animal procedures were approved by the Institutional Animal Care and Use Committee. Cortical neurons were isolated from embryonic day 17 (E17) C57BL/6J mouse embryos (Charles River) of mixed sex. Cortices were dissected in ice-cold HBSS, digested in 0.05% trypsin-EDTA (Thermo Fisher Scientific) for 10 min at 37°C, and triturated in MEM (Thermo Fisher) containing 0.6% glucose (Sigma) and 10% FBS (Hyclone). Cell viability was assessed with Trypan Blue (Sigma), and 50,000 viable neurons were spot-plated onto the center of poly-D-lysine–coated 22 mm coverslips (Matsunami Inc) to generate high-density networks. Cultures were maintained at 37°C, 5% CO₂, and 95% relative humidity in glial-conditioned Neurobasal Plus medium (Thermo Fisher) supplemented with GlutaMAX (Gibco) and B27 plus (Invitrogen). Half-medium exchanges were performed 2–3 times per week; no antibiotics or antimycotics were used. For plasmid-based co-transfection (TMEM106B constructs with pGW1-mApple), neurons were transfected at 4 days in vitro (DIV4) using Lipofectamine 2000 (Invitrogen) according to the manufacturer’s protocol. For AAV-mediated expression, neurons were transduced at DIV4 with AAV8-TMEM106B-FL, -CTF, or -gCTF (titer-matched) and, where applicable, co-transduced with AAV-PHP.eB-GFP-NLS.

### Longitudinal Microscopy

Primary cortical neurons co-transfected at DIV4 with pGW1-mApple and TMEM106B constructs (or empty vector) were imaged every 24 h for 10 days on a Keyence BZ-X810 microscope equipped with a ×10 objective and an environmental chamber maintaining 37°C and 5% CO₂. Stitched, focus-stacked images were analyzed in ImageJ. Neuronal death was scored blind to genotype and defined by soma rounding, neurite retraction, or loss of fluorescent signal on any given day, with the imaging timepoint assigned as the time of death. Cumulative survival curves were generated using the Kaplan–Meier method, and differences between conditions were assessed by Cox proportional hazards regression with empty-vector–transfected neurons as the reference group. Individual neurons (n ≥ 150 per condition, across 3 independent biological replicates) were tracked longitudinally.

### Immunofluorescence — cell lines

U2OS or SH-SY5Y cells grown on coverslips were fixed with 4% paraformaldehyde (Electron Microscopy Sciences) for 20 min at room temperature and washed three times in PBS (Corning). Cells were permeabilized in 0.2% saponin (Sigma) in PBS for 10 min, then blocked in 4% BSA (Sigma) in PBS for 30 min at room temperature. Primary antibodies were applied in blocking buffer overnight at 4°C. The following primary antibodies were used: mouse anti-HA (1:500; Biolegend #901501), mouse anti-LAMP1 (1:200; DSHB #H4A3-c), rabbit anti-TDP-43 (1:500; Proteintech #12892-1-AP), mouse anti-phospho-TDP-43 (1:500; Cosmo Bio #CAC-TIP-PTD-M01), rabbit anti-Lamin B1 (1:500; Proteintech #12987-1-AP), rabbit anti-LAP1 (1:200; Proteintech #21459-1-AP), rabbit anti-KPNB1 (1:200; Abcam #ab2811), and mouse anti-RanGAP1 (1:500; Santa Cruz #sc-28322). Amyloid-like inclusions were detected with Amytracker 540 (1 μM; Ebba Biotech) co-incubated with primary antibodies. After three PBS washes, species-matched Alexa Fluor-conjugated secondary antibodies (Invitrogen, 1:500) were applied for 1 h at room temperature in the dark. Coverslips were washed, counterstained with DAPI, and mounted with ProLong Gold Antifade (Invitrogen). Images were acquired on a Keyence BZ-X810 microscope with a ×60 oil immersion objective using matched acquisition parameters across conditions.

### Immunofluorescence — primary cortical neurons

Primary cortical neurons were fixed, permeabilized, and blocked as described for cell lines. The following primary antibodies were used: mouse anti-HA (1:500; Biolegend #901501), rabbit anti-TDP-43 (1:500; Proteintech #12892-1-AP), rabbit anti-Lamin B1 (1:500; Proteintech #12987-1-AP), rabbit anti-LAP1 (1:200; Proteintech #21459-1-AP), and guinea pig anti-MAP2 (1:500; Synaptic Systems #188-004). Secondary antibody incubation, mounting, and imaging were performed as above.

### Immunofluorescence - human postmortem tissue

Frontal cortex sections from neurologically normal individuals were provided by the Emory Alzheimer’s Disease Research Center and the VA Biorepository Brain Bank with Institutional Review Board approval. Case demographics, including age, sex, and postmortem interval, are summarized in Supplementary Table 1. Paraffin-embedded 6 μm sections were deparaffinized at 60°C for 15–20 min, cleared through three 15 min Histoclear (National Diagnostics) incubations, and rehydrated through graded ethanol (100%, 90%), distilled water, and PBS. Antigen retrieval was performed by incubating the sections in 80% formic acid for 10 min, followed by antigen retrieval in 10 mM citrate buffer (pH 6.0) at 120°C in a steamer for 25 min with a 30 min cool-down. After PBS rinses, tissue boundaries were marked with a hydrophobic pen, and sections were permeabilized with 0.2% saponin (Sigma) for 10 min. Blocking was performed for 1 h at room temperature in 1.5% normal donkey serum in PBS. Primary antibodies diluted in blocking solution were applied overnight at 4°C: rabbit anti-TMEM106B luminal domain (1:500; Synaptic Systems #506 017), guinea pig anti-NeuN (1:500; Synaptic Systems #266004), rabbit anti-TDP-43 (1:500; Proteintech #12892-1-AP), rabbit anti-Lamin B1 (1:500; Proteintech #12987-1-AP), rabbit anti-LAP1 (1:200; Proteintech #21459-1-AP), and mouse anti-TOM20 (1:50; Santa Cruz #sc-17764). Sections were washed and incubated with Alexa Fluor-conjugated secondary antibodies (Invitrogen, 1:500) in 1.5% normal donkey serum for 1 h at room temperature. Lipofuscin autofluorescence was quenched with TrueBlack (Biotium; diluted in 70% ethanol) for 30s. After final washes in PBS and distilled water, slides were coverslipped with ProLong Gold Antifade containing DAPI (Invitrogen) and cured overnight. Imaging was performed on a Keyence BZ-X810 with a ×60 oil immersion objective using matched acquisition settings across cases and analyzed in ImageJ.

### Image quantification

All quantitative image analysis was performed in ImageJ/FIJI by an investigator blinded to condition. Segmentation of nuclei was performed on DAPI-stained images using a combination of Gaussian blur, auto-thresholding (Otsu), and watershed separation. For TDP-43 nuclear-to-cytoplasmic (N/C) ratio quantification, nuclear masks were defined from DAPI and cytoplasmic masks from HA (transfected cells), MAP2 (primary neurons), or NeuN (human tissue) signal. Mean fluorescence intensity of TDP-43 within each compartment was background-corrected using a local non-cellular region, and the N/C ratio was calculated per cell. For GFP-NLS reporter quantification, the same nuclear/cytoplasmic segmentation strategy was applied. Nuclear circularity was calculated from DAPI masks using the ImageJ circularity metric (4π × area / perimeter²). Because Lamin B1 localizes to the nuclear envelope rather than the nucleoplasm, fluorescence intensity was quantified within a 3-pixel-wide band along the nuclear periphery, generated by dilating and subtracting the DAPI mask, as previously described^71^. For human tissue, NeuN-positive neurons were classified as TMEM106B fibril-positive (detectable punctate luminal-domain signal within the cell body) or fibril-negative by a blinded scorer; ambiguous cells were excluded. For each case, 50 fibril-positive and 55 fibril-negative neurons were quantified where available. LAMP1 fluorescence intensity and lysosomal area were quantified using auto-threshold segmentation of LAMP1 images followed by the Analyze Particles function. For colocalization with Amytracker 540, the Amytracker signal was thresholded and overlaid on the HA channel. All acquisition settings (laser power, exposure, gain) were held constant across conditions within each experiment.

### Protein lysate preparation and sequential fractionation

Transfected cells were harvested 48 h after post transfection, washed once with cold PBS, and lysed on ice in RIPA buffer (pH 7.4; Bio-world) supplemented with Halt protease and phosphatase inhibitor cocktail (Thermo Fisher). Lysates were sonicated (three 15 s pulses at 25% amplitude, separated by 5 s rest intervals) and clarified by centrifuge at 16,000 × g for 15 min at 4°C. The supernatant was collected as the RIPA-soluble fraction. The remaining pellet was washed three times with 1x PBS and resuspended in 8 M urea buffer (10 mM Tris, pH 8.0) with brief sonication to extract RIPA-insoluble, urea-soluble material. Protein concentrations in both fractions was determined by BCA assay (Pierce).

### Immunoblotting

Samples (20–30 μg per lane) were mixed with 4x Laemmli buffer and separated on 4–20% gradient SDS-PAGE gels (Bio-Rad). To preserve TMEM106B-FL glycosylated homodimers, those samples were loaded without heat denaturation; all other samples were denatured at 95°C for 5 min. Proteins were transferred onto PVDF membranes (Bio-Rad), blocked for 1 h at room temperature in 5% non-fat milk in PBS-T (0.1% Tween-20), and probed overnight at 4°C with the following primary antibodies diluted in blocking buffer: mouse anti-HA (1:1000; Biolegend #901501), rabbit anti-V5 (1:1000; Cell Signaling #13202), rabbit anti-APEX2 (1:1000; Cell Signaling #74728), mouse anti-actin (1:10,000; ABclonal #AC050), and rabbit anti-GAPDH (1:10,000; ABclonal #AC002). For N-linked deglycosylation, lysates were treated with PNGase F (New England Biolabs) according to the manufacturer’s protocol prior to SDS-PAGE. After washing in PBS-T, membranes were incubated for 1 h at room temperature with HRP-conjugated or IRDye-conjugated secondary antibodies (LI-COR). For detection of biotinylated proteins in APEX2 experiments, membranes were probed with IRDye 800CW streptavidin (LI-COR #926-32230). Chemiluminescent signal was detected using SuperSignal West Pico substrate (Pierce) and captured on a ChemiDoc imaging system (Bio-Rad).

### Co-immunoprecipitation

U2OS cells were seeded in 10 cm dishes and co-transfected with HA-tagged TMEM106B-FL or -gCTF (or empty vector) together with V5-tagged LAP1/TOR1AIP1 using polyethylenimine. Forty-eight hours after transfection, cells were washed with cold PBS and lysed in IP buffer (50 mM Tris-HCl pH 7.4, 150 mM NaCl, 1% Triton X-100, 1 mM EDTA) supplemented with Halt protease and phosphatase inhibitor cocktail (Thermo Fisher). Lysates were clarified by centrifugation at 16,000 × g for 15 min at 4°C, and protein concentration was determined by BCA assay. Equal amounts of protein (500 μg) were incubated with anti-HA magnetic beads (Thermo Fisher #88836) for 2 h at 4°C with rotation. A 5% aliquot was reserved as input. Beads were washed five times with IP buffer, and bound proteins were eluted by boiling in 2× Laemmli buffer for 5 min at 95°C. Inputs and immunoprecipitates were resolved by SDS-PAGE and immunoblotted with anti-V5, anti-HA, and anti-actin antibodies as described above

### APEX2 proximity labeling

SH-SY5Y cells were plated on poly-L-lysine-coated 10 cm dishes and transfected with TMEM106B-gCTF-V5-APEX2, V5-APEX2 (cytosolic control), or left untransfected. Each condition was performed in biological triplicates. Forty-eight hours after transfection, culture media was replaced with fresh media containing 500 μM biotin-phenol, and cells were incubated for 30 min at 37°C. Labeling was initiated by adding H₂O₂ to a final concentration of 1 mM and gently agitating for 1 min. Control plates received PBS in place of H₂O₂. The reaction was quenched by three rapid washes with quenching buffer (PBS containing 10 mM sodium azide, 10 mM sodium ascorbate, and 5 mM Trolox), followed by four 5 min incubations in quenching buffer supplemented with protease inhibitors on ice. Cells were lysed in RIPA-like buffer (50 mM Tris, 150 mM NaCl, 0.4% SDS, 0.5% sodium deoxycholate, 1% Triton X-100, 10 mM sodium azide, 10 mM sodium ascorbate, 5 mM Trolox, and protease inhibitor cocktail). Lysates were sonicated on ice (2 s on, 2 s off, 25% amplitude, 1 min total, repeated twice) and clarified by centrifugation at 16,500 × g for 10 min at 4°C. Protein concentration was measured by RC/DC assay (Bio-Rad).

### Streptavidin pulldown and mass spectrometry

Biotinylated proteins were enriched using Nanolink streptavidin magnetic beads. Beads were pre-washed three times in TBS containing 0.1% Tween-20, then incubated with 100 μg of clarified lysate overnight at 4°C with rotation. Beads were washed sequentially with RIPA buffer containing 0.4% SDS, 2% SDS in 50 mM Tris-HCl (pH 7.4), and two additional rounds of RIPA buffer with 0.4% SDS. A 10% aliquot of beads was reserved for streptavidin-blot validation. Remaining beads were washed four times in PBS and submitted to the Emory Integrated Proteomics Core (RRID:SCR_023530) for on-bead digestion and LC-MS/MS analysis as previously described^72^. Briefly, peptides were reconstituted in loading buffer (0.1% trifluoroacetic acid) and 2 μL of each sample was injected onto a self-packed C18 column (1.9 μm resin, Dr. Maisch, Germany; 15 cm × 100 μm ID; New Objective) coupled to an EASY-nLC 1200 system (Thermo Fisher Scientific). Separation was achieved over a 56 min gradient at 700 nL/min using mobile phase A (0.1% formic acid in water) and mobile phase B (0.1% formic acid in acetonitrile), ramping from 1% to 40% B over 56 min, then to 99% B within 1 min and holding for 3 min. Data were acquired on a Q-Exactive Plus mass spectrometer (Thermo Fisher Scientific) operating in data-dependent mode, collecting one full MS scan (400–1600 m/z, 1 × 10⁶ AGC, 100 ms maximum injection time, 70,000 resolution at m/z 200) followed by 20 HCD MS/MS scans (2 m/z isolation window, 28% normalized collision energy, 1 × 10⁵ AGC, 50 ms maximum injection time, 17,500 resolution at m/z 200). Previously sampled precursors were dynamically excluded for 20 s within a 10 ppm mass window; charge states of +1 and ≥+7 were excluded.

Raw files were searched against the human UniProt/Swiss-Prot database using the Andromeda search engine within MaxQuant. Variable modifications included methionine oxidation (+15.9949 Da), asparagine and glutamine deamidation (+0.9840 Da), and protein N-terminal acetylation (+42.0106 Da), with up to five modifications per peptide. Cysteine carbamidomethylation (+57.0215 Da) was set as a fixed modification. Fully tryptic peptides with up to two missed cleavages were considered, with a precursor mass tolerance of ±20 ppm prior to calibration and ±4.5 ppm after MaxQuant internal calibration. Additional parameters included a maximum peptide mass of 6,000 Da, minimum peptide length of six residues, and 0.05 Da tolerance for MS/MS scans. Peptide spectral match, protein, and site-level false discovery rates were each controlled at 1%. For label-free quantification, the MaxLFQ algorithm was applied using razor and unique peptides. Full MS1 features were matched across runs using a 0.7 min retention time window following alignment within a 20 min search space. Downstream analysis and visualization, including principal component analysis and volcano plots, were carried out in Perseus. Proteins with a fold change ≥ 2 and p < 0.05 were considered significantly enriched or depleted. Gene ontology (GO) enrichment analysis was performed using the clusterProfiler package in R, with significantly enriched proteins as input and the human proteome as background. Terms with adjusted p-value < 0.05 (Benjamini-Hochberg correction) were considered significant.

### Statistical analysis

All statistical tests and graphing were conducted using GraphPad Prism (version 10). Data are presented as mean ± SD. Comparisons between two groups were assessed by Student’s t-test, and comparisons among three or more groups were evaluated by one-way ANOVA followed by appropriate post hoc testing as indicated in figure legends. Neuronal survival data were analyzed by Kaplan-Meier estimation and compared using Cox proportional hazards regression. Data are presented as mean ± SD or mean ± SEM as indicated in figure legends. Sample sizes (n) refer to biologically independent replicates as defined for each experiment in the corresponding figure legend. A p-value < 0.05 was considered statistically significant (ns, not significant; *p < 0.05; **p < 0.01; ***p < 0.001; ****p < 0.0001).

## Data and code availability

All data supporting the findings of this study are available from the corresponding author upon reasonable request.

## Contributions

Conceptualization: KT, JJ. Methodology: KT, JP, MD, DP, FM, JZ, TB. Investigation: KT, JP, MD, DP, FM, JZ, JJ. Visualization: KT, JP, MD, DP. Resources: JJ. Funding acquisition: KT, JJ. Project administration: JJ. Supervision: JP, JJ. Writing–original draft: KT, JJ. Writing–review and editing: KT, JP, JJ.

## Acknowledgements

We would like to thank Dr. Jonathan Glass and Dr. Schniederjan from Emory University, and Dr. Ian Robey from the VA Biorepository for providing access to the clinical materials. We would also like to thank Dr. Gary Bassell, Dr. Zachary McEachin, Dr. Thomas Kukar and their lab members for the discussion and support. KT was supported by an R36 dissertation fellowship (R36AG088283). JP is supported by AAN/AFTD/American Brain Foundation (0000082888). JJ is supported by the National Institute of Health (R01AG068247 and R01NS138605). This work was supported in part by the Emory Integrated Proteomics Core (RRID:SCR_023530).

