## Supplemental Figures for "TMEM106B C-terminal fragments drive nucleocytoplasmic transport failure and TDP-43 mislocalization in the aging human brain"

**Supplementary Figure 1. TMEM106B fibrils co-label with Amytracker 540 in aged human frontal cortex and full-length TMEM106B selectively expands the lysosomal compartment.**

**(A)** Representative immunofluorescence images from the frontal cortex of a 72-year-old neurologically normal individual, stained with Amytracker 540 (green), anti-TMEM106B luminal domain (TMEM-CTF; magenta), and DAPI (blue). **(B)** Representative immunofluorescence images of U2OS cells expressing HA-tagged empty vector (CTRL), TMEM106B-FL, CTF, or gCTF, stained for HA (green), LAMP1 (red), and DAPI (blue). Arrows (FL row) indicate markedly enlarged LAMP1-positive lysosomes; arrowhead (gCTF row) highlights a representative transfected cell. **(C)** Quantification of LAMP1 integrated density per cell across conditions. Violin plots show median (solid line) and interquartile range (dashed lines). Statistics: one-way ANOVA with Tukey's post-hoc test; n =50 cells per condition across 3 independent experiments. Scale bar, 10 μm.

**Supplementary Figure 2. TMEM106B fragments drive TDP-43 mislocalization in U2OS cells without inducing phosphorylated TDP-43 inclusions.**

**(A)** Representative immunofluorescence images of U2OS cells expressing HA-tagged C-terminal 25 kDa TDP-43 fragment (TDP-25; positive control), TMEM106B-FL, CTF, or gCTF, stained for HA (green), phosphorylated TDP-43 (pTDP-43, Ser409/410; magenta), and DAPI (blue). **(B)** Representative immunofluorescence images of U2OS cells expressing HA-tagged empty vector (CTRL), TMEM106B-FL, CTF, or gCTF, stained for HA (green), total TDP-43 (red), and DAPI (blue). Dashed outlines demarcate nuclei of transfected cells. TMEM106B-CTF and gCTF drove redistribution of endogenous TDP-43 from the nucleus to the cytoplasm, whereas CTRL and TMEM106B-FL cells maintained nuclear TDP-43 enrichment. Scale bar, 10 μm. **(C)** Quantification of TDP-43 N/C ratio across conditions (corresponding to panel B). CTF and gCTF significantly reduced the TDP-43 N/C ratio compared with CTRL, whereas FL did not differ significantly from CTRL. Violin plots show median and interquartile range. *$p < 0.05$; ns, not significant (one-way ANOVA with Tukey's post-hoc test). Scale bar, 10 μm.

**Supplementary Figure 3. TMEM106B fibril-positive neurons show reduced TDP-43 nuclear localization across additional aged individuals, while a young fibril-negative case retains normal TDP-43 distribution.**

**(A)** Representative immunofluorescence images of additional aged human frontal cortex cases (Cases #3, #4, #5, and #6) stained with DAPI and anti-TMEM106B luminal domain (TMEM-CTF; cyan/blue), TDP-43 (red), and NeuN (green). Filled arrowheads indicate fibril-positive neurons; open arrowheads indicate fibril-negative neurons. **(B)** Representative immunofluorescence images from a 22-year-old neurologically normal individual (Case #7), stained for the same markers. NeuN+ neurons show intact nuclear TDP-43 localization and no detectable TMEM-CTF signal, consistent with the age-dependent accumulation of TMEM106B fibrils. Scale bar, 10 μm.

**Supplementary Figure 4. TMEM106B CTF and gCTF expression disrupt nuclear shape, nuclear lamina organization, and the nuclear import machinery in U2OS cells.**

**(A)** Representative immunofluorescence images of U2OS cells expressing HA-tagged empty vector (CTRL), TMEM106B-FL, CTF, or gCTF, stained for HA (green) and DAPI (blue, shown in merged panel). Dashed outlines delineate nuclei;

arrowheads indicate nuclear indentations in CTF-expressing cells. **(B)** Quantification of nuclear circularity (4π × area / perimeter²) in U2OS cells. Violin plots show median and interquartile range. **(C)** Representative immunofluorescence images of primary cortical neurons transduced with AAV encoding CTRL, TMEM106B-FL, CTF, or gCTF, stained for HA (green) and DAPI (blue). MAP2 is shown in the merged panel (magenta). **(D)** Quantification of nuclear circularity in primary cortical neurons. **(E)** Representative immunofluorescence images of U2OS cells expressing the same constructs, stained for HA (green), Lamin B1 (red), and DAPI (blue). Arrowheads indicate reduced Lamin B1 nuclear rim intensity in CTF- and gCTF-expressing cells. **(F)** Quantification of Lamin B1 mean intensity at the nuclear rim in U2OS cells. **(G)** Representative immunofluorescence images of U2OS cells stained for HA (red), KPNB1/Importin-β1 (green), and DAPI (blue). Arrowheads indicate diminished perinuclear KPNB1 enrichment and aberrant cytoplasmic distribution in CTF- and gCTF-expressing cells.. **(H)** Representative immunofluorescence images of U2OS cells stained for HA (red), RanGAP1 (green), and DAPI (blue). Arrowheads indicate disrupted RanGAP1 distribution in CTF- and gCTF-expressing cells. ***$p < 0.001$; **$p < 0.01$; *$p < 0.05$; ns, not significant (one-way ANOVA with Tukey's post-hoc test). Scale bar, 10 μm.

**Supplementary Figure 5. Gene Ontology and subcellular compartment enrichment analyses of the gCTF-APEX2 proximal proteome.**

Gene Ontology (GO) Biological Process enrichment dot plot for proteins significantly enriched in the gCTF-APEX2 proximal proteome. Top enriched terms include protein folding, response to ER stress, the ERAD pathway, glycoprotein metabolic process, protein folding in ER, response to unfolded protein, and ER-to-Golgi vesicle transport, consistent with gCTF's luminal biogenesis route. Nucleocytoplasmic transport and nuclear pore complex assembly also emerged as significantly enriched terms. Dot size reflects gene count; color scale indicates $-\log_{10}$ adjusted p-value.

**Supplementary Figure 6. Lamin B1 and LAP1 disorganization in TMEM106B fibril-positive neurons across additional aged cases and absence of these phenotypes in a young control.**

**(A)** Representative immunofluorescence images of postmortem frontal cortex from four additional aged, neurologically normal individuals (Cases #3, #4, #5, #6) stained for TMEM-CTF/DAPI (cyan/blue), Lamin B1 (red), and NeuN (green). Fibril-positive NeuN+ neurons display fragmented or discontinuous Lamin B1 signal (filled arrowheads) compared with intact rim staining in neighboring fibril-negative neurons (open arrowheads). **(B)** Representative immunofluorescence images of the same additional aged cases stained for TMEM-CTF/DAPI (cyan/blue), LAP1 (red), and NeuN (green) **(C)** Representative immunofluorescence images from the young control individual (Case #7, age 22) stained for TMEM-CTF/DAPI (cyan/blue), Lamin B1 (top row, red), LAP1 (bottom row, red), and NeuN (green). Scale bar, 10 μm.

**Supplementary Figure 7. Cytoplasmic mitochondrial signal is preserved in TMEM106B fibril-positive neurons with disrupted Lamin B1**

Representative immunofluorescence images of postmortem human frontal cortex (Case #1) showing a fibril-negative neuron (TMEM−, top) and a fibril-positive neuron (TMEM+, bottom) from the same tissue section, stained for TMEM-CTF/DAPI (cyan/blue), Lamin B1 (red), and TOM20 (green, mitochondrial marker), with merged overlay. Scale bar, 10 μm.

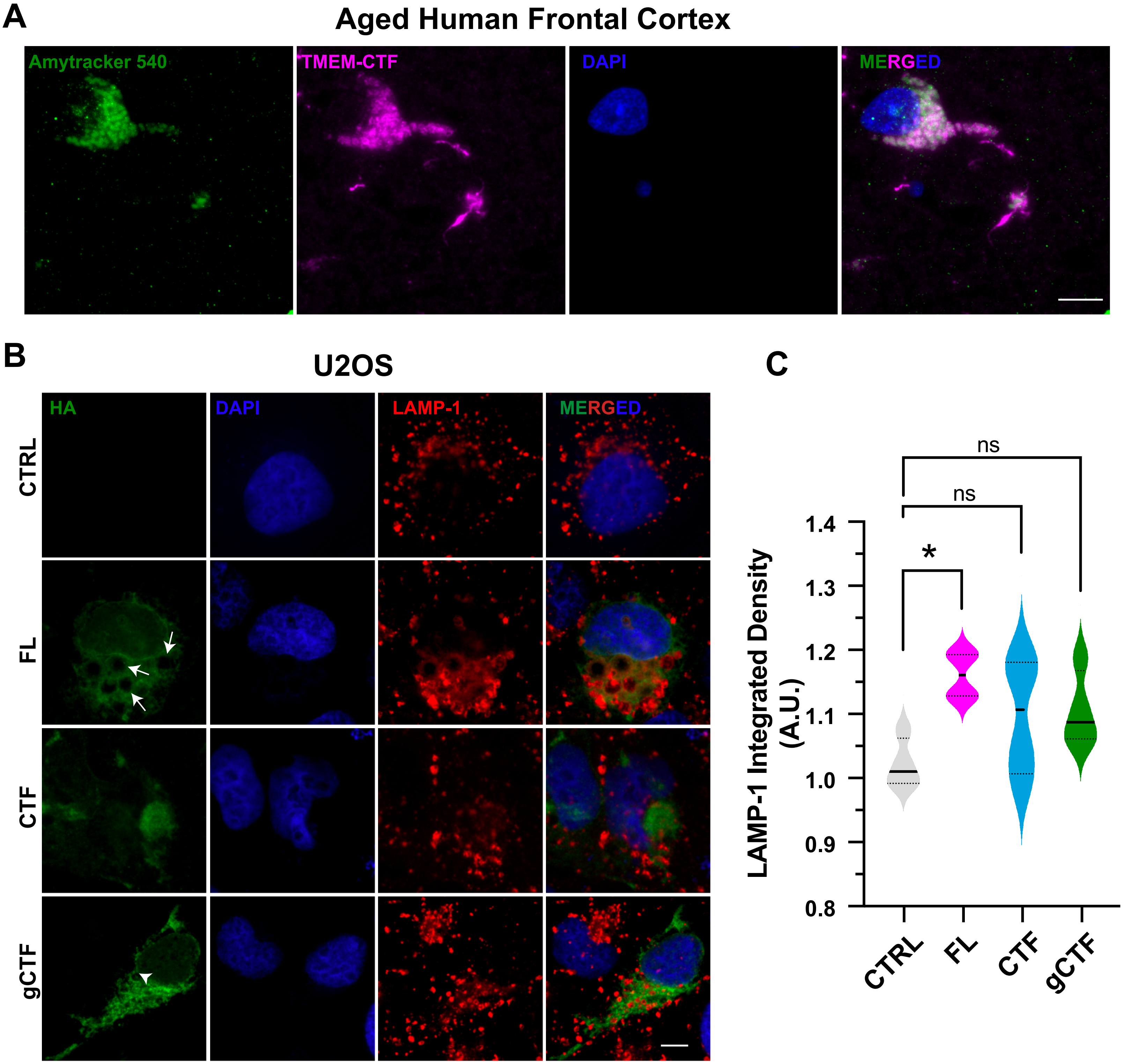


Supplementary Figure 1

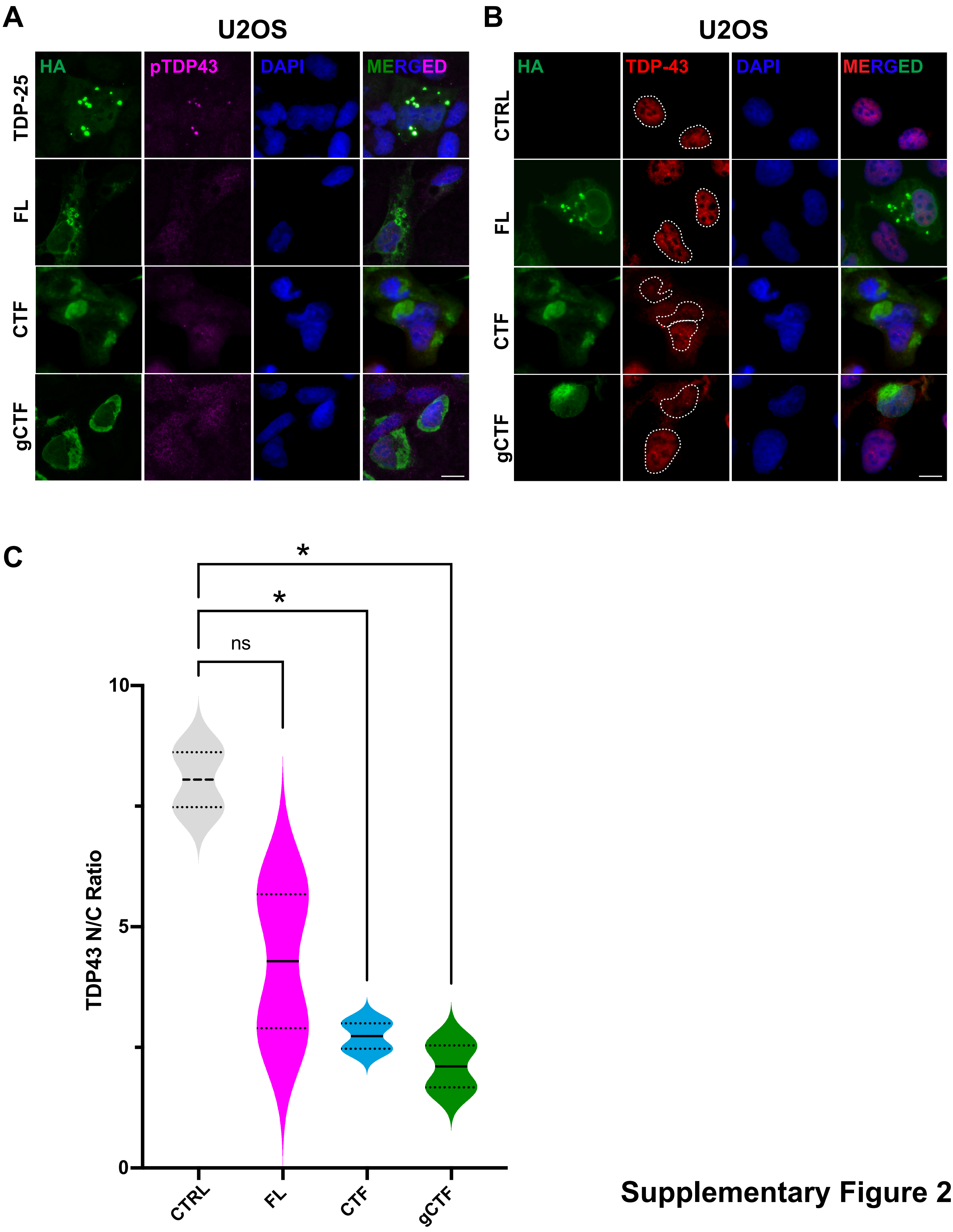


Supplementary Figure 2

A

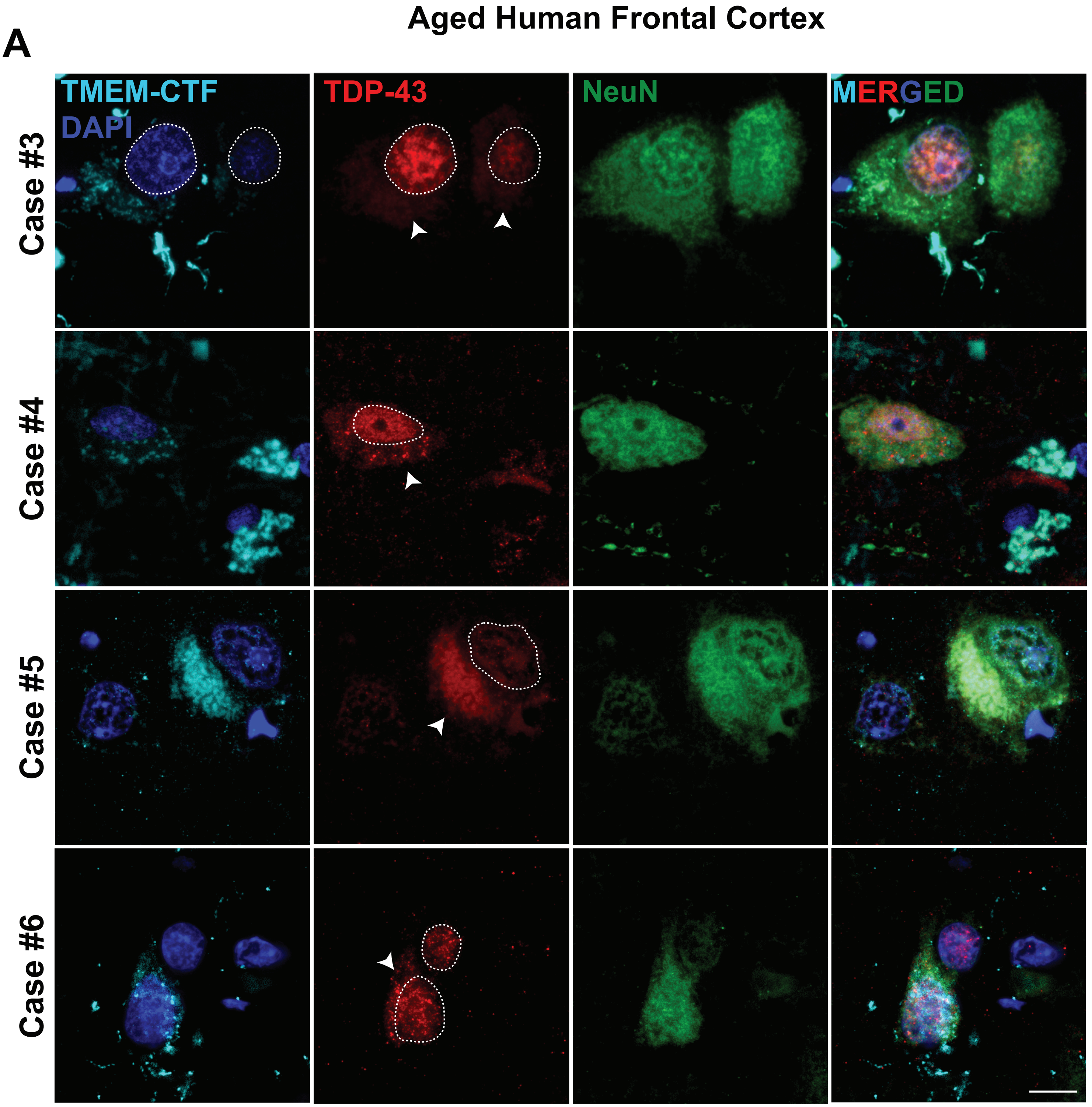


B

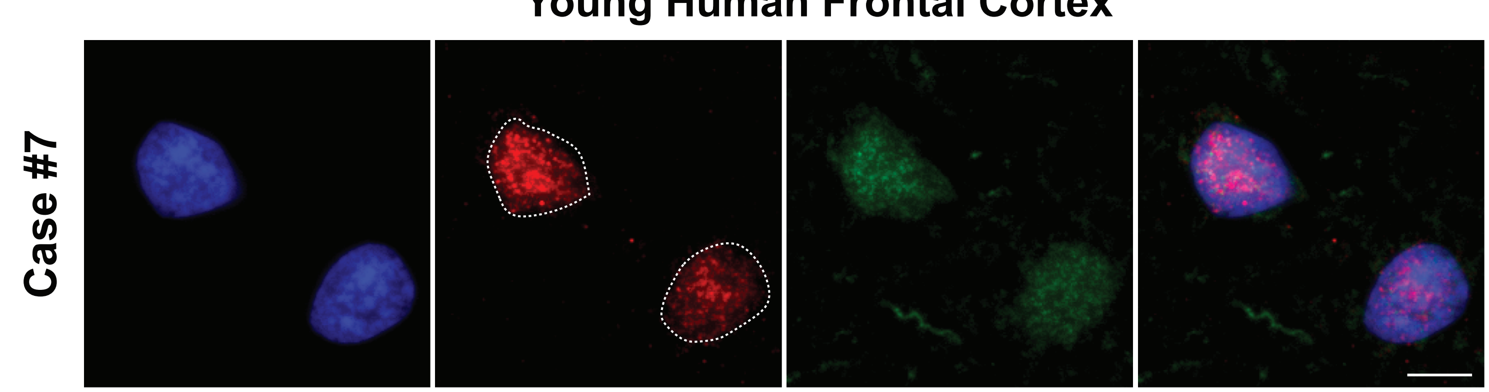


Supplementary Figure 3

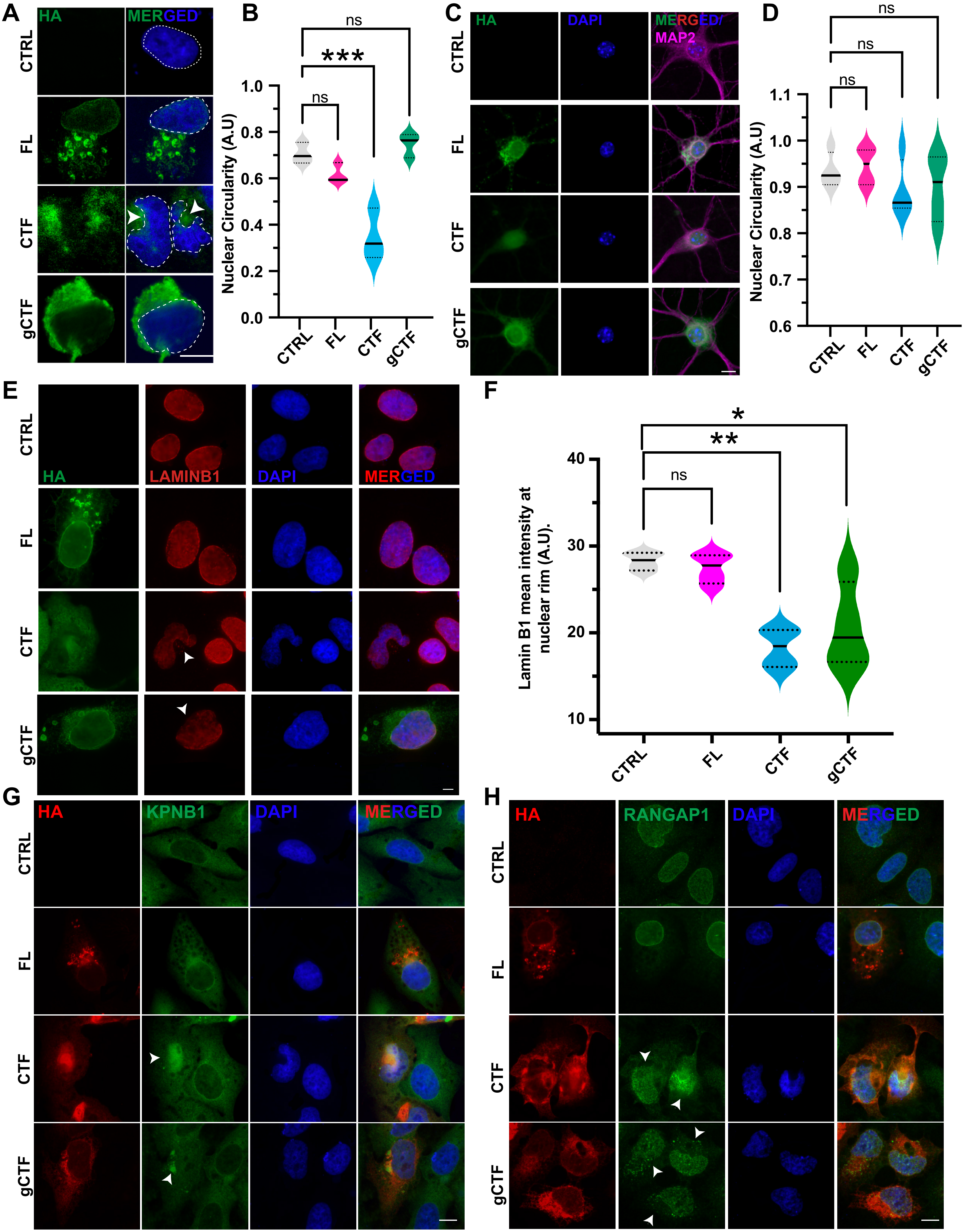


### Supplementary Figure 4

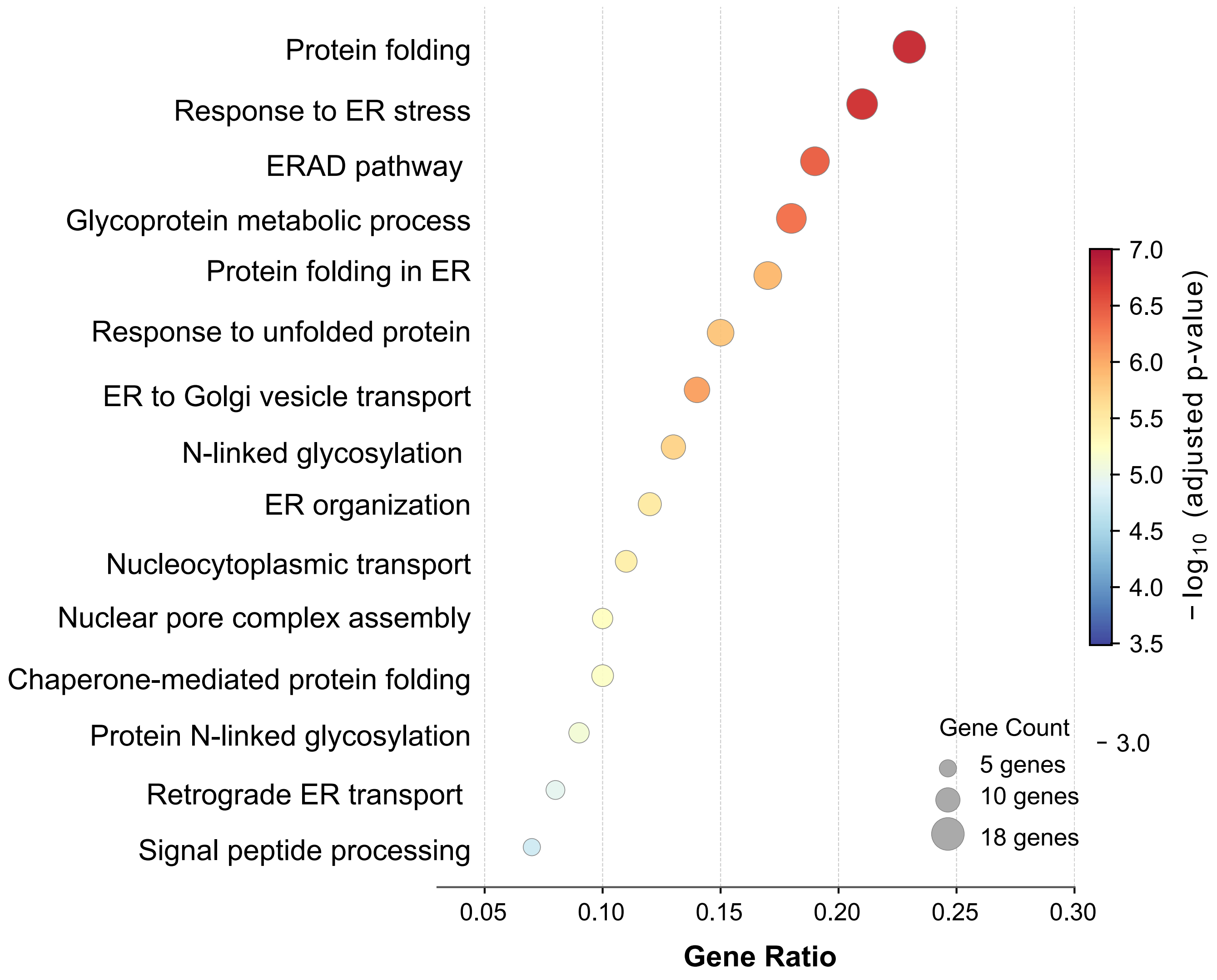


**Supplementary Figure 5**

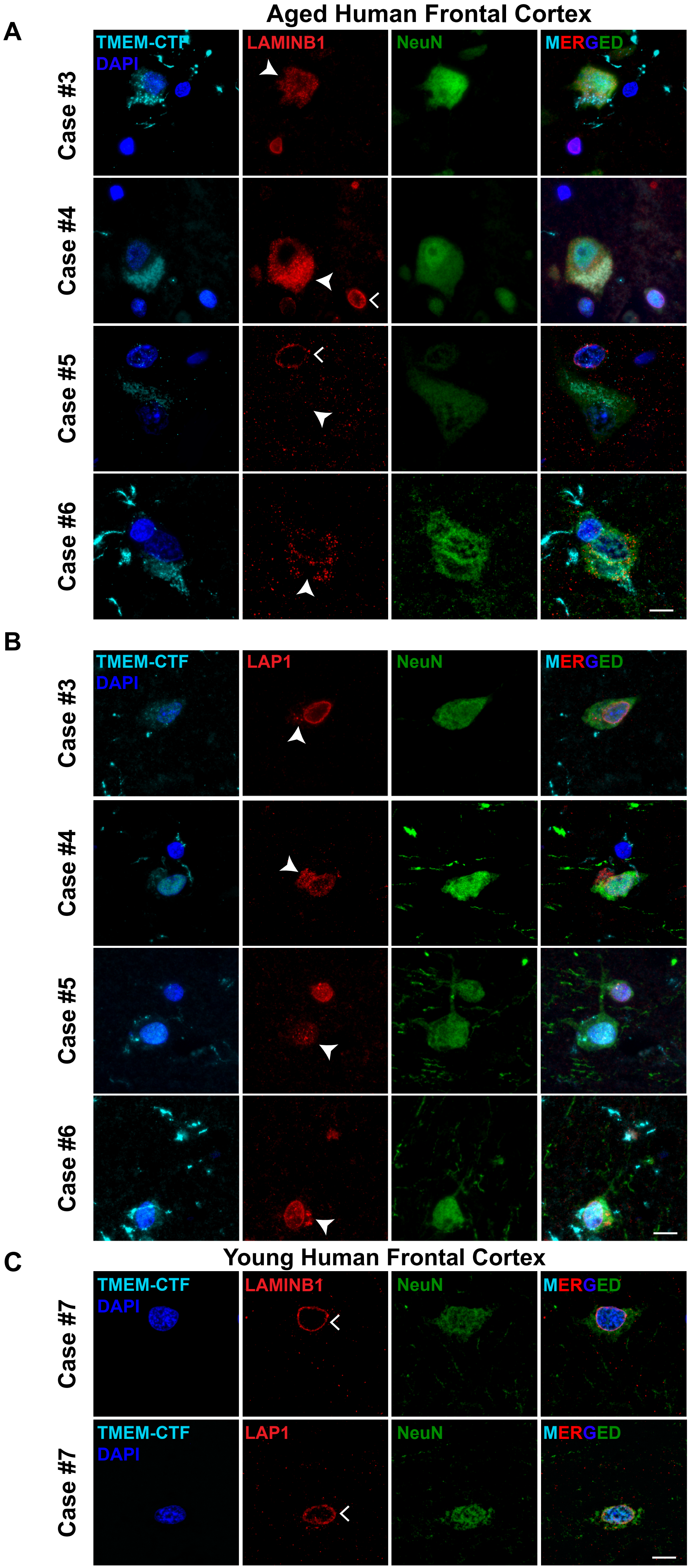


Supplementary Figure 6

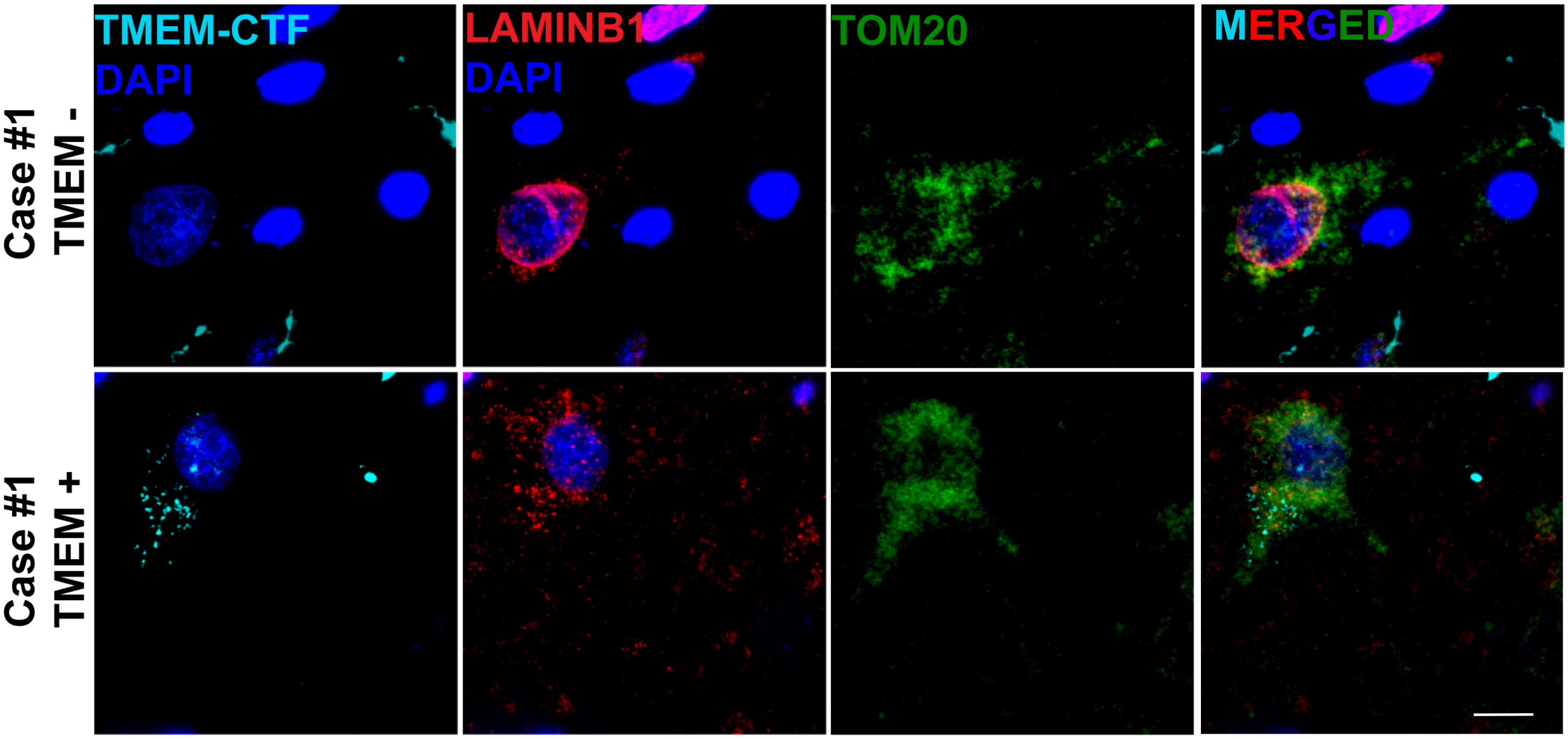


**Supplementary Figure 7**
