## Supplemental Table 1 for "TMEM106B C-terminal fragments drive nucleocytoplasmic transport failure and TDP-43 mislocalization in the aging human brain"

### Supplementary Table 1. Human brain tissue donors used in this study.

| Case ID | Sex | Age (yr) | Ethnicity | PMI-frz (hr) |
|---|---|---|---|---|
| **Case 1** | M | 78 | W | 27.4 |
| **Case 2** | M | 84 | W | 75.0 |
| **Case 3** | M | 70 | W | 26.7 |
| **Case 4** | M | 72 | W | 112.75 |
| **Case 5** | M | 74 | W | 7 |
| **Case 6** | M | 69 | W | 6 |
| **Case 7** | M | 22 | W | |

*All cases are neurologically normal aged controls obtained from the VA Biorepository Brain Bank (Southern Arizona VA).*

*Abbreviations: PMI-frz, post-mortem interval to freezing (hours); W, White/Caucasian;*
